# Endocannabinoids facilitate transitory reward engagement through retrograde gain-control

**DOI:** 10.1101/2025.01.06.630792

**Authors:** David J. Marcus, Anthony E. English, Gunn Chun, Emmaline F. Seth, Rachel Oomen, Sabrina Hwang, Bailey Wells, Sean C. Piantadosi, Azra Suko, Yulong Li, Larry S. Zweifel, Benjamin B. Land, Nephi Stella, Michael R. Bruchas

## Abstract

Neuromodulatory signaling is poised to serve as a neural mechanism for gain control, acting as a crucial tuning factor to influence neuronal activity by dynamically shaping excitatory and inhibitory fast neurotransmission. The endocannabinoid (eCB) signaling system, the most widely expressed neuromodulatory system in the mammalian brain, is known to filter excitatory and inhibitory inputs through retrograde, pre-synaptic action. However, whether eCBs exert retrograde gain control to ultimately facilitate reward-seeking behaviors in freely moving mammals is not established. Using a suite of in vivo physiological, imaging, genetic and machine learning-based approaches, we report a fundamental role for eCBs in controlling behavioral engagement in reward-seeking behavior through a defined thalamo-striatal circuit.

## 1 INTRODUCTION

Reward-seeking behaviors are evolutionarily conserved and necessary for survival, and importantly disrupted in both substance-use disorder and numerous psychiatric diseases ^1,2,3,4,5^. The Nucleus Accumbens (NAc) is a key nodal mesocorticolimbic structure that mediates action selection and as well as reward-seeking ^6,7,8,9,10^. The NAc incorporates input from numerous excitatory, inhibitory, and neuromodulatory projections to ultimately integrate affective information and translate this information into motivated behavioral outputs ^11,12,13^. However, we lack a complete understanding of critical fundamental players in reward-seeking, specifically whether diverse sources of excitatory input can be integrated and filtered by distinct neuromodulators to shape discrete behavioral outputs in the NAc. For example, the NAc receives glutamatergic input from over half a dozen structures, yet it is unclear why such diverse sources of this single neurotransmitter exist and how each input is filtered into useful information downstream by local modulatory signaling ^14,15,16^.

Brain slice electrophysiological studies have shown that he eCB signaling system dynamically regulates both excitatory and modulatory input into the NAc ^17,18,19,20,21^. This system, composed principally of the cannabinoid 1 and cannabinoid 2 (CB_1_R and CB_2_R) receptors as well as the ligands anandamide and 2-arachidonylglycerol (2-AG), represents the most widely expressed neuromodulatory system in the mammalian brain ^22,23,24,25,26^. Unlike conventional neurotransmitters released by presynaptic vesicular mechanisms, lipophilic eCBs are produced on-demand by post-synaptic neurons and primarily function retrogradely to activate presynaptic CB_1_R to attenuate neurotransmitter release ^27,28,29,30^. Furthermore, the mechanistic link between neural activity-dependent eCB release, modulation and shaping of afferent input, and whether specific behavioral features are elicited by this retrograde action *in vivo* remain to be established. Here we discovered specific behavioral parameters necessary to evoke eCB release in the NAc *in vivo* during freely moving motivated behaviors. Our results indicate that eCB production and release is time locked to behavioral engagement through a genetically defined medium spiny NAc neuron population interacting with presynaptic paraventricular thalamic inputs. This retrograde signaling dynamically shapes the vigor of engagement in reward seeking behaviors through discrete suppression of extrinsic glutamatergic inputs into the NAc.

## 2 RESULTS

### Endogenous cannabinoids are released in the NAc during reward consumption to inhibit presynaptic glutamate release

To investigate how eCB dynamics in the NAc facilitate reward seeking and consumption, we injected the medial NAc shell of adult mice with an AAV to drive the neuronal expression of the fluorescent eCB biosensor, GRAB_eCB2.0_, and implanted photometry fibers dorsal to the injection site (Fig 1A) ^31^. Mice were then placed in a Pavlovian reward conditioning assay, in which mice were trained to associate a cue light with the protraction of a sipper containing a 10% sucrose solution across 5 days. On day 1 of training, consumption of the sucrose solution elicited a pronounced increase in GRABeCB2.0 fluorescence, indicative of local eCB release time-locked to consummatory events (Fig 1B). However, on day 5 of conditioning, the onset of the increase in GRAB_eCB2.0_ fluorescence shifted leftward toward the time of cue onset, and rose to 50% of its maximal response prior to sipper delivery. This suggests that eCB release is not solely tied to reward consumption, but may sculpt reward-seeking behaviors (Fig 1C-E, S1 A-G). Accordingly, in a reward omission test in which the sipper was not delivered on 50% of cue trials, the GRAB_eCB2.0_ signal still increased despite a lack of reward delivery (Figure S1H,I. To determine which eCB ligand responsible for the GRAB_eCB2.0_ signal, we treated mice with vehicle or DO34 (20mg/kg, i.p), an inhibitor of the rate limiting enzyme in 2-AG biosynthesis (Diacyl-glycerol Lipase, DAGL). DO34 strongly attenuated the reward-induced increase in GRAB_eCB2.0_ signal, suggesting that 2-AG is mediating this response. As a selectivity control, we confirmed that the CB_1_R antagonist, SR141716 (10 mg/kg, i.p.) blocked this increase in GRAB_eCB2.0_ signal (Fig S1J-O). To determine whether eCB release in the NAc is selective for stimulus valence or unique to reward-seeking behavior, we exposed the same cohorts of animals to a Pavlovian fear conditioning assay. We found that shock exposure also elicited elicited a significant increase in GRAB_eCB2.0_ signal, suggesting that eCBs production in the NAc is elicited by both salient rewarding and aversive stimuli (Fig S1P,Q). However, during fear retrieval, exposure to the shock-predictive tone did not elicit any increase in the GRAB_eCB2.0_ signal (Fig S1R). Together, these results indicate that both rewarding and aversive stimuli can elicit robust eCB release within the NAc (Fig S1S).

**FIGURE 1.**
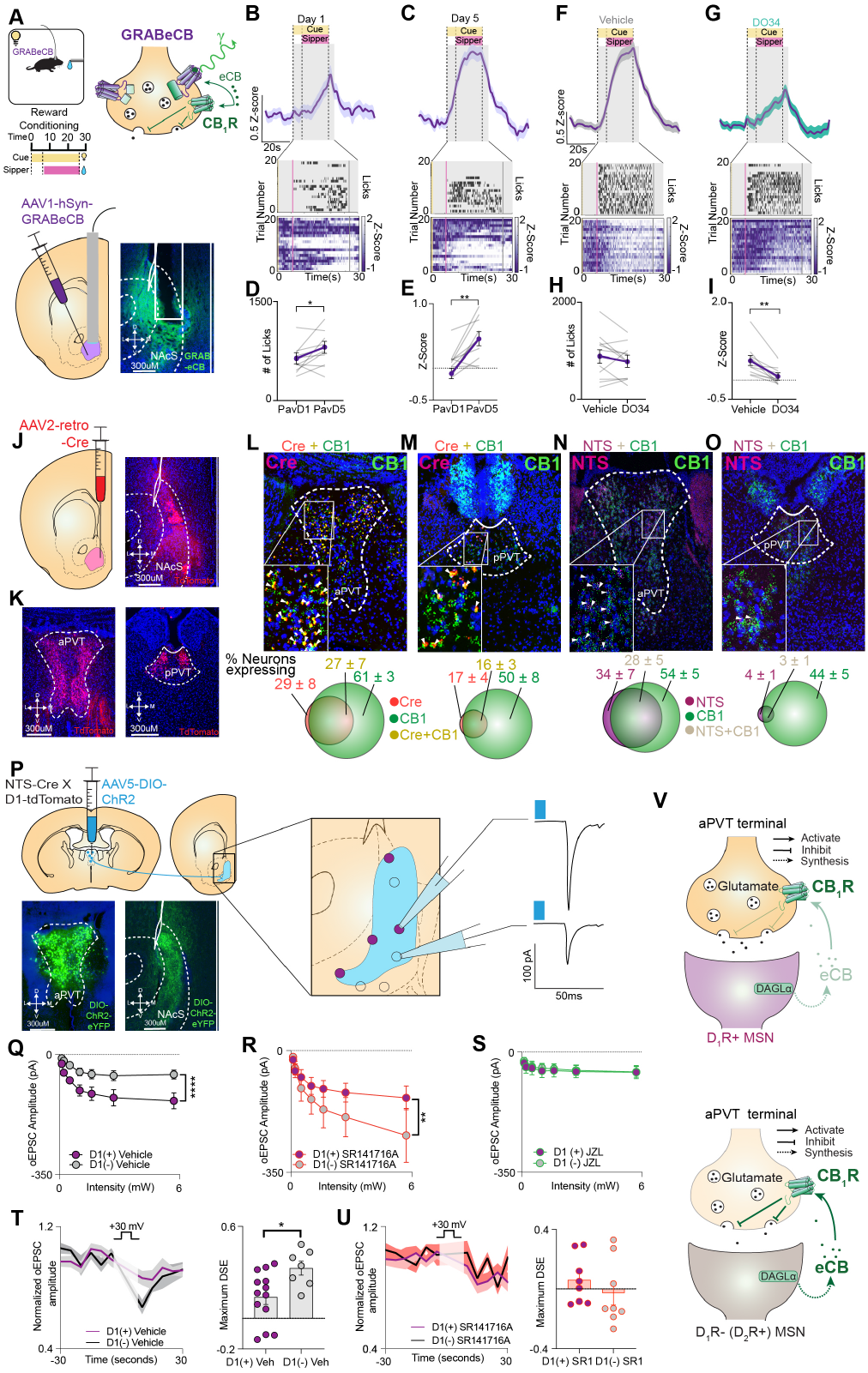
Endogenous cannabinoids are released in the NAc during reward consumption and inhibit presynaptic glutamate release. Schematic for fiber photometry recordings of GRAB_eCB2.0_ in the NAc during Pavlovian reward conditioning (A). Photometry trace with representative lick raster and heatmap of GRAB_eCB2.0_ signal aligned to cue onset on day 1 (B) and day 5 (C) of conditioning. Comparison # of licks per session (p=0.0468)(D) and average photometry Z-score from 0-26s (p=0.0027)(E) between day 1 and day 5 (N=11). Trace with representative lick raster and heatmap of GRAB_eCB2.0_ signal aligned to cue onset after treatment with vehicle (F) or DO34 (G). Comparison # of licks per session (p=0.2514)(H) and average photometry Z-score from 0-26s (p=0.0011) (I) between vehicle and DO34 (N=11). Schematic for retrograde tracing and RNAscope studies in Ai14 mice (J). Epifluorescent images of TdTomato positive NAc projecting neurons in the aPVT and pPVT (K). RNAscope image and quantification of Cre and CB_1_R transcript levels in the aPVT (L) and pPVT (M). RNAscope image and quantification of NTS and CB_1_R transcript levels in the aPVT (N) and pPVT (O). Schematic for whole-cell voltage-clamp recordings of optically evoked EPSCs from NAc D_1_R+ and D_1_R-neurons (P). Optically evoked input-output curve from D_1_R+ and D_1_R-neurons in vehicle (D_1_R+ n=19, D_1_R-n=20, p<0.001) (Q) SR141617A (D_1_R+ n=14 D_1_R-n=12, p=0.0050) (R) and JZL184 (D_1_R+ n=17, D_1_R-n=15, p=0.9698) (S) treated slices. Recordings of depolarization-induced suppression of excitation in vehicle (D_1_R+ n=13, D_1_R-n=7, p=0.0184) (T) and SR141716A (D_1_R+ n=8, D_1_R-n=8, p=0.3763) (U) treated slices. Summary schematic depicting enhanced retrograde eCB signaling at aPVT^NTS^-NAc D_1_R+ synapses compared to aPVT^NTS^-NAc D_1_R-synapses (V). All error bars represent ± SEM; N represents number of mice, n represent number of neurons. p values reported from two-tailed paired t test (D,E,H,I), two-tailed unpaired t test (T,U), and two-way ANOVA (Q,R,S). \**p* < 0.05, \*\**p* < 0.01, \*\*\**p* < 0.001, \*\*\*\**p* < 0.0001.

Considering that a primary function of 2-AG activation of presynaptic CB_1_R is to modulate fast neurotransmitter release, we next identified neuronal projections to the NAc that express CB_1_R (Fig 1J) ^27^. We injected a retrograde virus to drive the expression of Cre recombinase (AAV2-retro-hSyn-Cre) into the NAc of flox-stop Ai14 Td-Tomato reporter mice, and identified five principal sources of excitatory inputs: the medial prefrontal cortex (mPFC), anterior paraventricular thalamus (aPVT), posterior paraventricular thalamus (pPVT), basolateral amygdala (BLA), and ventral hippocampus (vHipp) (Fig 1K, S2A-C). We then used a quantitative form of in situ hybridization, RNAscope, to examine the colocalization of Cre mRNA transcript (NAc projectors) and *cn1r* mRNA transcript. Overall, the aPVT had the largest percentage of neurons projecting to the NAc, as well as the largest percentage of neurons projecting to the NAc that coexpressed CB_1_R (Fig 1L.M, S2 D-F).

Recent studies have emphasized the functional differences that exist along the anterior-posterior axis of the PVT, yet few genetic markers have been identified to isolate and manipulate neurons located in the anterior paraventricular thalamus (aPVT) ^32,33,34,35,36,37,38,39^. We analyzed the Allen Brain Atlas database and published PVT RNAseq results and found that the expression of neuropeptide neurotensin (NTS) is selectively enriched in the aPVT compared to the posterior paraventricular thalamus (pPVT) ^40,41^. We further corroborated these initial findings with RNAscope, showing selective expression in the aPVT. We also found that in aPVT NTS neurons showed strong coexpression of CB_1_R (83.9 ± 5%)(Fig 1 N,O). To examine how aPVT input to the NAc is regulated by CB_1_R signaling, we performed whole-cell electrophysiological recordings and Channelrhodopsin (ChR2)-assisted circuit mapping ^42^. NTS-Cre mice were crossed with a D_1_R-tdTomato reporter mice, and their progeny and were injected with AAV5-DIO-ChR2-eYFP, facilitating recordings of optically-evoked excitatory postsynaptic currents (oEPSCs) from either D_1_R+ are D_1_R-(purportedly D_2_R+, see methods and Fig S2G-N) neurons in the medial NAc shell (Fig 1P). Photo-stimulation induced input/output curves demonstrate that there is a significant bias in excitatory input to D_1_R+ neurons compared to D_1_R-(Fig 1Q). Remarkably treatment with SR141716A inverted this bias, resulting in larger larger excitatory responses observed in D_1_R-compared to D_1_R+ (Fig 1R). In agreement with the involvement of 2-AG in this response, the intrinsic excitatory bias onto D_1_R+ neurons could be eliminated by augmenting 2-AG levels by inhibiting its degradation by Monoacylglycerol Lipase using its the inhibitor, JZL 184 (Fig 1S).To determine if a phasic 2-AG signaling is involved, we recorded depolarization-induced suppression of excitation (DSE), a form of short-term synaptic plasticity response known to be mediated by retrograde 2-AG signaling. We found that DSE was selectively expressed at aPVT-NAc D_1_R-synapses (Fig T,U). These data suggest that both tonic and phasic eCB signaling are primarily expressed at aPVT-NAc D_1_R-synapses, and that tonic eCB signaling at these synapses drives the observed excitatory bias onto D_1_R+ neurons (Fig 1V). Taken together, these *in vivo* and in vitro results demonstrate that salient rewarding and aversive stimuli evoke robust eCB production and that 2-AG release modulates extrinsic excitatory input to the NAc using discrete cell type-specific retrograde signaling.

### aPVT-NAc terminal activity is negatively correlated with engagement in reward-seeking behaviors

To assess how eCB signaling facilitates reward-seeking behaviors through filtering of extrinsic excitatory input, we first determined the behavioral role of the aPVT^NTS^-NAc circuit using a suite of *in vivo* optical approaches to record from and manipulate the activity of these projections. First, we injected NTS-Cre mice with AAV-DJ-GCaMP6s in the aPVT followed by fiberoptic implantation in the medial NAc shell to assess calcium dynamics of aPVT^NTS^-NAc terminals during consummatory and reward-seeking behaviors (Fig 2A). During ad libitum sucrose access, we observed a significant decrease in GCaMP6s fluorescence time-locked to the first lick in a lick bout, as previously reported ^14,15,32,33^(Fig 2B). Interestingly, when the GCaMP6s signal was aligned to the last lick in a bout, there was a noted increase in aPVT^NTS^-NAc activity (Fig 2C). These findings indicate that the aPVT^NTS^-NAc circuit may not solely encode information relevant to the stimulus valence or salience, but instead may shape and refine the degree of animal’s approach or “behavioral engagement.” To further assess the role of aPVT^NTS^-NAc circuit in regulating behavioral engagement, mice subsequently underwent Pavlovian reward conditioning. Here, the GCaMP6s signal was aligned to the reward-predictive cue and significant differences emerged across conditioning days (Fig 2D,E, Fig S3A-M). Key changes in calcium activity were isolated during the cue period prior to sipper protraction (0-6 seconds), and in the period immediately follow cue cessation and sipper retraction (26-46 seconds). Specifically, in the pre-sipper time window (0-6s), we found a transition from primarily excitation on day 1 to primarily inhibition on day 5 (Fig 2F). Conversely, in the time window following cue offset and sipper retraction (26-46s), we observed a transition from primarily inhibition on day 1 to excitation on day 5 (Fig 2G). Analyzing videos (1080p, 60 hz) of mice behaviors throughout the sessions revealed substantial behavioral changes as the mice learned the task parameters. *In particular, we observed interesting and significant changes in engagement with the sipper port, which we quantified as % of time poking, pawing, or sniffing at the sipper port* (Sup video S1 and S2, methods). On day 1 of conditioning, mice only spent roughly 1/4 of their time (24.1%) engaging with the sipper port during the cue window prior to sipper protraction (0-6s), as they had not yet learned to associate the cue with the rewarding outcome. However, by day 5 of conditioning, mice spent 2/5 (38.6%) of their session time poking, pawing, and sniffing at the sipper port during the cue window, in a display of learned expectation of reward delivery (Fig 2H,). Conversely, on day 1 of conditioning in the post-sipper time window (26-46s), mice spent almost 1/3 (31.8%) of the time engaging with the sipper port following sipper retraction, as they had not yet learned the associated time-out period (ITI) following sipper retraction. However, by day 5 of conditioning, this behavior had significantly diminished to less than 1/5 (19.0%) of the time, as the mouse learned that the sipper will not be extended again until the next cue delivery (Fig 2I, Sup video S2).(Fig 2I, Sup video S2). We found that whether computing either the pre-sipper or post-sipper time window, the activity of the aPVT^NTS^-NAc circuit was significantly negatively correlated with an animals relative epochs of sipper port engagement, suggesting that this signals engagement and disengagement from reward-seeking. (Fig 2J,K).

**FIGURE 2.**
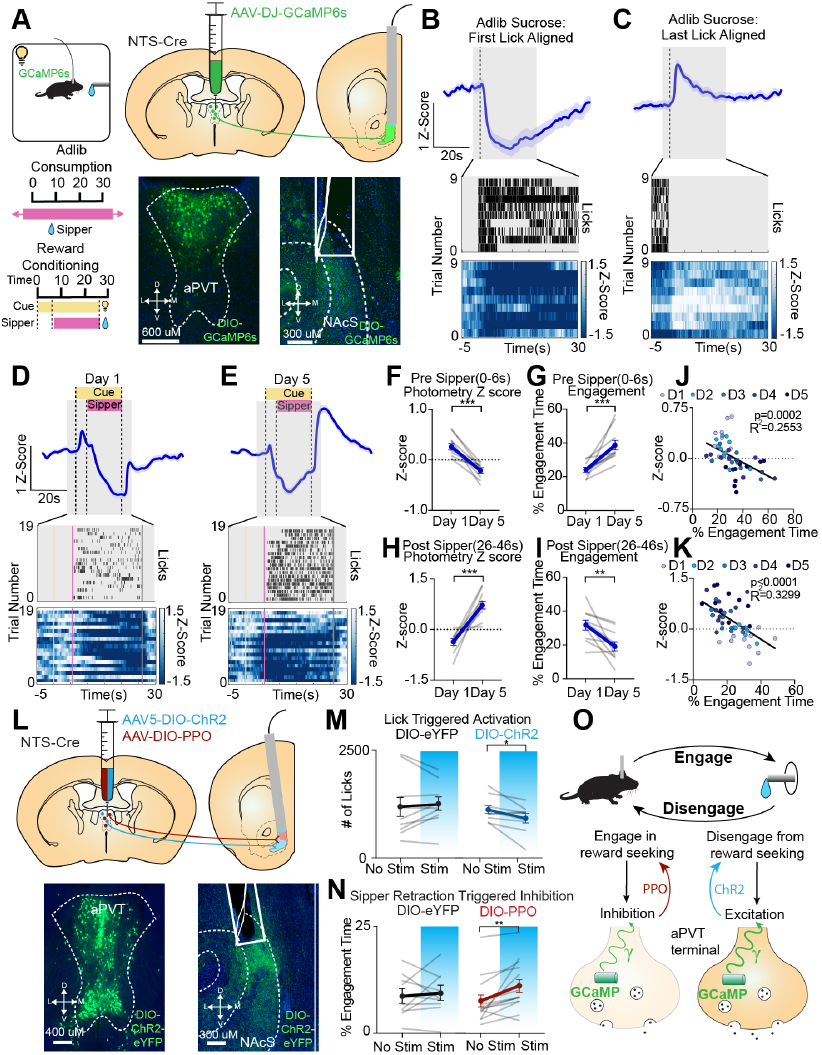
aPVT-NAc terminal activity is negatively correlated with engagement in reward-seeking behaviors. Schematic for fiber photometry recordings of GCaMP6s expressing aPVT^NTS^-NAc terminals (A). Photometry trace with representative lick raster and heatmap of first lick (B) and last lick (C) aligned GCaMP6s signals during adlib consumption. Photometry trace with representative lick raster and heatmap of cue-aligned GCaMP6s signal during day 1 (D) and day 5 (E) of Pavlovian reward conditioning (N=10). Comparison between day 1 and day 5 of average photometry Z-score (p=0.0001) (F) and sipper port engagement time (p=0.0002) (G) during the six second cue window prior to sipper extension. Correlation between pre-sipper extension engagement time and average photometry Z-score across 5 days of conditioning (R^2^=0.2553, p=0.0002) (H). Comparison between day and day 5 of average photometry Z-score (p=0.0001) (I) and sipper port engagement time (p=0.0039) (J) during the twenty second time window following sipper retraction. Correlation between post-sipper retraction engagement time and average photometry Z-score across 5 days of conditioning (R^2^=0.3299, p=0<0.0001) (K). Schematic for optogenetic manipulation of aPVT-NAc terminals using DIO-ChR2 and DIO-PPO (L). # of licks during no stimulation or closed loop 20hz optogenetic stimulation of aPVT^NTS^-NAc terminals in DIO-ChR2 (N=9, p=0.0300) and DIO-eYFP (N=9, p=0.6323) expressing animals (M). # of licks during no stimulation or sipper retraction-aligned 10hz optogenetic inhibition of aPVT-NAc terminals in DIO-PPO (N=13, p=0.0090) and DIO-eYFP (N=11, p=0.7608) expressing animals (N). Summary schematic depicting behavioral engagement-induced inhibition of aPVT^NTS^-NAc terminals (O). All error bars represent ± SEM; N represents number of mice. p values reported from two-tailed paired t test (F,G,H,I,M,N) and linear regression (J,K) \**p* < 0.05, \*\**p* < 0.01, \*\*\**p* < 0.001.

To assess whether aPVT^NTS^-NAc terminal inhibition measured during the sipper protraction was driven solely by reward consumption events, we used a reward omission experiment. In rewarded trials, calcium dynamics mirrored the dynamics observed on day 5 of reward conditioning (inhibited during cue/reward, excited post cue/reward) (Fig S4A-C). In the omitted trials, aPVT terminals were similarly inhibited despite a lack of reward delivery, suggesting that these inhibitory events are not specifically relevant for reward consumption (Fig S2C). Another possibility is that incentive salience had been attributed to the cue, and that the reward-predictive cue had become an attractive/reinforcing target. To specifically examine this, trials from day 5 of Pavlovian reward conditioning were isolated in which the animals ignored sipper delivery events. We reasoned that if aPVT^NTS^-NAc activity encodes the incentive salient features of the cue, aPVT terminals should be inhibited regardless of whether the animal engages the sipper port. We found that the aPVT^NTS^-NAc terminals are not inhibited during ignored trials, indicative of activity of the circuit being tied to the patterns and features of behavioral engagement (Fig S4D). Lastly, to determine whether the activity of this circuit is simply due to any locomotor behavioral event (inhibited during decrease, excited during increase), we conditioned mice in an FR1 task where reward delivery requires an action-outcome motor sequence (poke, walk to sipper port, consume). We found that even during engagement in active motor behaviors to obtain reward, aPVT^NTS^-NAc was was broadly inhibited, suggesting its activity is not specifically tied to locomotion (Fig S4E-H). This was further corroborated by a test of open field activity, in which we observed a lack of calcium dynamics time-locked to locomotor initiation or rearing (Fig S4I-K).

To determine whether activity of the aPVT^NTS^-NAc terminals was only correlated with engagement in reward-seeking behaviors, we trained the mice in an aversive Pavlovian fear conditioning and extinction task (Fig S5A). During fear conditioning we observed a transient (5 second) peak in GCaMP6s signal time-locked to foot shock (Fig S5B). On day 1 of fear extinction (recall), we observed a significant inhibition of aPVT^NTS^-NAc terminals, similar to what we observed during engagement in reward seeking behaviors, again suggesting that this circuit does not signal stimulus valence (Fig S5C). When the photometry signal was aligned to the end of freezing bouts (disengagement), there was a concomitant increase in fluorescence, again suggesting this circuit broadly signals behavioral engagement (Fig S5D). As mice underwent extinction training, the inhibitory signal broadly decreased, and their degree of engagement in defensive freezing behaviors was likewise negatively correlated with the activity of the aPVT^NTS^-NAc terminals(Fig S5 E-H). These results collectively establish that dynamic aPVT^NTS^-NAc activity is negatively correlated with engagement in both reward-seeking and defensive freezing behavior.

We next examined whether we could selectively modulate behavioral engagement by photo-manipulating the neuronal activity of the aPVT^NTS^-NAc circuit (Fig 2L) during different reward-seeking epochs. Mice were injected with either AAV5-DIO-ChR2-eYFP, AAV5-DIO-PPO-eYFP (an inhibitory Gi-coupled opsin), or AAV5-DIO-YFP as control in the aPVT and bilateral fiber optic implants were placed above the medial NAc shell. Following Pavlovian reward conditioning, mice injected with ChR2 or eYFP received counterbalanced days of closed-loop optogenetic stimulation, in which each lick resulted in a 2 second burst of 20hz blue light photo-stimulation delivered to aPVT^NTS^-NAc terminals. Stimulation of aPVT^NTS^-NAc terminals resulted in reduced licking behavior, either using closed-loop photo-stimulation or constant 20hz photo-stimulation throughout the duration of the cue (Fig 2M,N, Fig S6A,B). Conversely, 10hz optogenetic Gi-mediated photo-inhibition of this circuit, from 5 seconds before sipper retraction to 20 seconds after, significantly increased engagement with the sipper port (Fig 2O,P). To test whether activation of the aPVT^NTS^-NAc circuit is necessary and sufficient for conditioned freezing behaviors, we either selectively activated or inhibited terminals using ChR2 or PPO respectively throughout the cue period (20hz 20 seconds and 10hz 20 seconds, respectively) during fear retrieval. Photo-activation of the aPVT terminals was sufficient to reduce conditioned freezing, while photo-inhibition resulted in a trend towards increased freezing, likely due to a ceiling effect on the expression of freezing behavior during recall (Fig S5I,J). Importantly, neither activation nor inhibition of the aPVT^NTS^-NAc circuit was innately reinforcing or aversive, as these photo-manipulations had no effect on valence in either intracranial self-stimulation of real-time place preference paradigms (Fig S6E-H). Lastly, photo-activation of the aPVT^NTS^-NAc terminals did not cause broad changes in locomotor behavior (Fig S3C,D). Overall, these findings establish the necessity and sufficiency of aPVT^NTS^-NAc projections in dynamic tuning of behavioral engagement in both an appetitive and aversive behavior (Fig 2Q).

### Endogenous cannabinoid inhibition of aPVT terminals drives engagement in reward-seeking behaviors

Our electrophysiology results revealed that 2-AG inhibits aPVT^NTS^-NAc terminals by activating CB_1_R and our *in vivo* calcium recordings indicate broad inhibition of activity of aPVT^NTS^-NAc terminals during epochs of behavioral engagement. Thus we tested whether this *in vivo* inhibition was mediated by endogenous 2-AG–CB_1_R signaling. Using mice expressing a DIO-GCaMP6s in aPVT^NTS^-NAc terminals, we found that systemic administration of the CB1 antagonist SR141716A(10mg/kg) blocked *in vivo* inhibition of these terminals, as well as significantly reduced sucrose consumption events (Fig 3A-E). Given that SR141716A was systemically delivered as is known to have anhedonic qualities, it remained unclear where the SR141716A was acting on CB_1_Rs to elicit these effects. Therefore, we next used a CRISPR/Cas9 viral approach to selectively delete CB_1_R from aPVT neurons to determine whether 2-AG–CB_1_R inhibition of aPVT^NTS^-NAc terminals was necessary for engagement in reward-seeking behaviors. NTS-Cre mice were injected with a single vector CRISPR/Cas9 virus which carried a guide RNA directed against the against a sequence in the first coding exon of the CB1 receptor (sgCNR1) or a guide RNA directed against the ROSA26 locus (sgROSA) to control for double stranded breaks (Fig 3F and Fig S7) ^43^. Fluorescent *in situ* hybridization showed that the sgCNR1 virus elicited a 47% reduction in the number of aPVT neurons coexpressing NTS and CB_1_R (30.4% vs 16.2%) compared to sgROSA26 controls (Fig S7A-H). To assess how deletion of aPVT CB_1_R affected aPVT^NTS^-NAc terminal activity as well as engagement in reward-seeking behaviors, we co-injected these viruses with AAV5-DIO-GCaMP6s in the aPVT, and implanted a fiber optic above the medial NAc shell. While this deletion did not alter the total number of licks, it significantly reduced the length of licking bouts (Fig S8A-G). This suggests that the CB_1_R receptor on aPVT neurons controls the innate temporal patterns of reward engagement, rather than impacting satiety-related behavioral measures.

**FIGURE 3.**
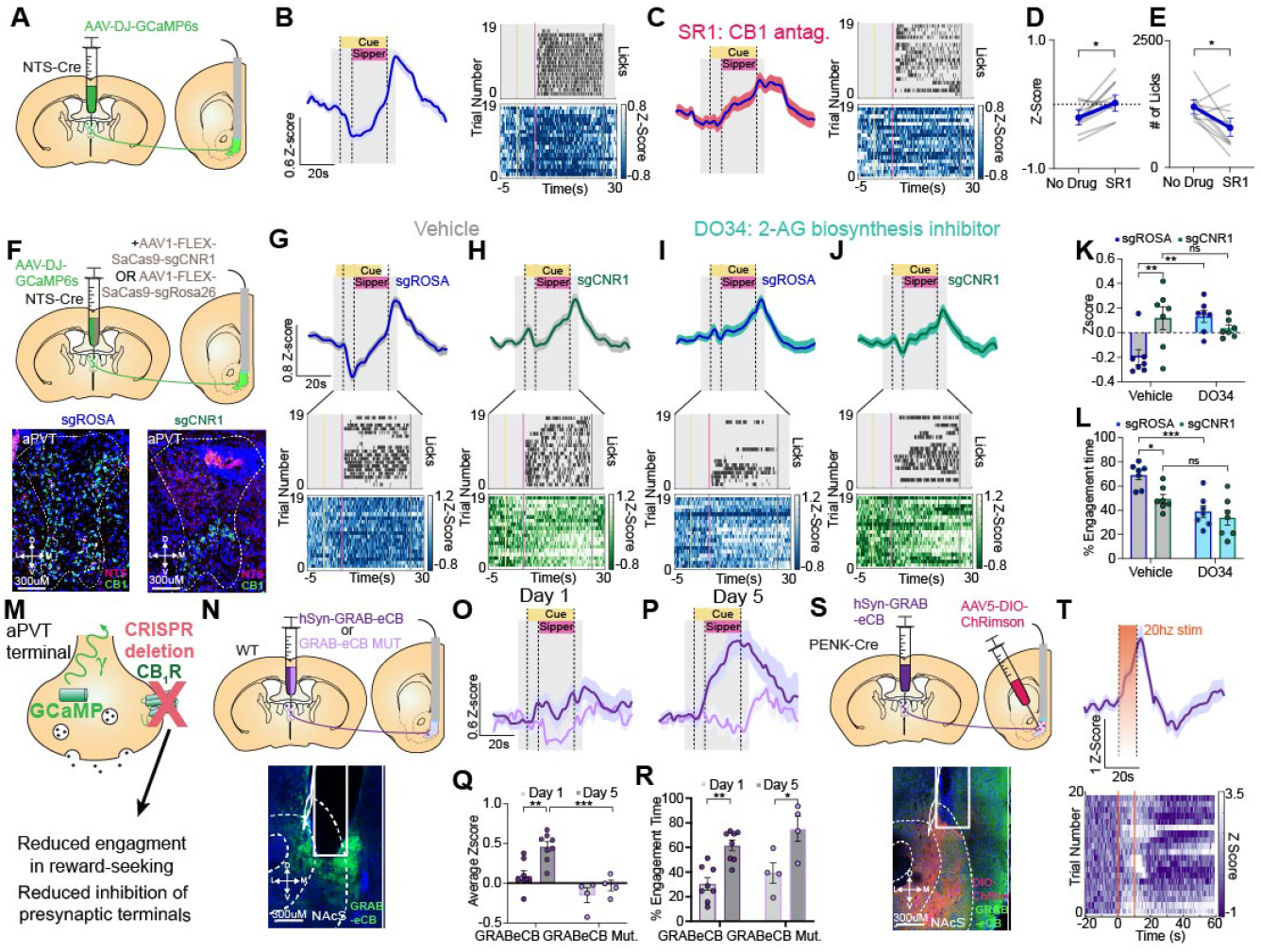
Endogenous cannabinoid inhibition of aPVT terminals drives engagement in reward-seeking behaviors. Schematic for photometry recordings of GCaMP6s expressing aPVT^NTS^-NAc terminals during Pavlovian reward conditioning (A). Photometry trace with representative lick raster and heatmap of GCaMP6s signal from no drug (B) and SR141716A (C) treated mice (N=10). Comparison of average Z-score from 0-26s (p=0.0064)(D) and # of licks per session (p=0.0270) (E) between vehicle and SR141716A treated mice. Schematic for CRISPR-Cas9 deletion of CB_1_R from aPVT NTS neurons and photometry recordings of aPVT^NTS^-NAc terminals during Pavlovian reward conditioning (F). Photometry trace with representative lick raster and heatmap of GCaMP6s signal from Vehicle (G,H) and DO34 (I,J) treated sgROSA (N=7) and sgCNR1(N=7) injected mice. Comparison of average photometry Z-score from 0-26 seconds showing blunted inhibition of aPVT terminals in vehicle treated sgCNR1 animals (p=0.0064) and DO34 treated sgROSA animals (p=0.0052) compared to vehicle treated sgROSA controls (K). Comparison of % sipper port engagement time showing reduced engagement in vehicle treated sgCNR1 animals (p=0.0293) and DO34 treated sgROSA animals (p=0.0008) compared to vehicle treated sgROSA controls (L). Summary schematic depicting beahvioral and physiological consequences of knocking out CB_1_R from aPVT^NTS^neurons (M). Schematic for photometry recordings of GRAB_eCB2.0_ or GRAB_eCB2.0MUT_ in aPVT-NAc terminals during Pavlovian reward conditioning (N). Photometry trace of GRAB_eCB2.0_ (N=8) or GRAB_eCB2.0MUT_ (N=4) signal on day 1 (O) and day 5 (P) of conditioning. Comparison of average photometry Z-score showing enhanced GRAB_eCB2.0_ signal of day 5 of reward conditioning compared to GRAB_eCB2.0_ day 1 (p=0.0017) and GRAB_eCB2.0MUT_ day 5 (p=0.0009) (Q). Comparison of % sipper port engagement time showing enhanced engagement time on day 5 in both GRAB_eCB2.0_ (p=0.0024) and GRAB_eCB2.0MUT_ (p=0.0120) compared to their respective day 1 values (R). Schematic for simultaneous optogenetic activation of NAc Penk neurons and photometry recordings of aPVT-NAc GRAB_eCB2.0_ (S). Trace and heatmap showing GRAB_eCB2.0_ dynamics time-locked to 10s 20hz stimulation of NAc Penk neurons (T). All error bars represent ± SEM; N represents number of mice. p values reported from two-tailed paired t test (D,E) and two-way ANOVA (K,L,Q,R) \**p* < 0.05, \*\**p* < 0.01, \*\*\**p* < 0.001.

To assess the pharmacological mechanism mediating the reward-engagement-induced inhibition of aPVT^NTS^-NAc terminals as well as the behavioral consequences of CB_1_R deletion from aPVT neurons, we treated GCaMP6s/sgCNR1 and GCaMP6s/sgROSA expressing mice with DO34 (inhibitor of 2-AG biosynthesis, 20mg/kg) or a vehicle control. In vehicle treated animals, CB_1_R deletion from aPVT neurons significantly attenuated the engagement-induced inhibition of aPVT^NTS^-NAc terminals, as well significantly reduced time engaging with the sipper port throughout the cue period (Fig 3G,H,K,L, S8H-K). This change was not due to a reduction in the number of engagement events, since the absence of engagement-related inhibition was still present in sgCNR1 when specifically isolating trials where animals consumed sucrose (Fig S8L). In DO34 treated animals, CB_1_R deletion had no effect either on aPVT^NTS^-NAc GCaMP6 signal or degree of engagement in reward-seeking (Fig 3I,J,K,L, S8D,E). Furthermore, DO34 reversed engagement-induced inhibition of aPVT^NTS^-NAc terminals in sgROSA mice and reduced sipper engagement, while having no effect on GCaMP6s signal or sipper engagement in sgCNR1 knockout animals (Fig 3K,L). Given that our histological characterization of CRISPR knock-down revealed a 50 % reduction in NTS neurons expressing CB_1_R, we tested whether increasing 2-AG signaling with JZL184 (an inhibitor of 2-AG hydrolysis) was sufficient to restore engagement-induced inhibition of aPVT^NTS^-NAc terminals to rescue the reward engagement deficits in sgCNR1 mice. JZL184 restored engagement-induced inhibition of aPVT^NTS^-NAc terminals and rescued engagement deficits in sgCNR1 mice to equivalent levels observed in controls (Fig S8O-R). Lastly, in sgCNR1 mice, there was no statistical correlation between behavioral engagement and aPVT-NAc terminal activity, suggesting decoupling of a key physiological mechanism regulating reward engagement (Fig S8S,T). The mutual occlusion between sgCNR1 knockdown and DO34 treatment provide strong corroborating evidence that 2-AG-CB_1_R signaling drives engagement in reward-seeking through inhibition of aPVT^NTS^-NAc terminals. These results also indicate that 2-AG-CB_1_R signaling represents a key endogenous Gi-GPCR mechanism which acts to tune excitatory input during engagement epochs of reward-seeking (Fig 3L).

We next assessed whether CRISPR knockdown of aPVT CB_1_R impaired engagement in defensive freezing behaviors in a Pavlovian fear conditioning/extinction paradigm (Fig S9A). During fear conditioning, deletion of the aPVT CB_1_R neither affected the activity of aPVT^NTS^-NAc terminals nor freezing behaviors (Fig S9B,C,F,G). Similarly, during fear recall, this manipulation did not attenuate freezing-induced inhibition of aPVT^NTS^-NAc terminals or freezing behaviors (Fig S9D,E,H,I). These data suggest that while inhibition of aPVT^NTS^-NAc terminals represents a common feature of engagement in both reward-seeking and freezing behaviors, 2-AG – CB_1_R signaling at aPVT^NTS^-NAc terminals is necessary for engagement in reward-seeking but not so for freezing behaviors, suggesting a unique mechanism driving aversion-related inhibition.

To determine whether 2-AG binds to presynaptic CB_1_R on aPVT terminals in the NAc, we injected mice with AAV1-hSyn-GRAB_eCB2.0_ or the control virus mutant-GRAB_eCB2.0_ (GRAB_eCB2.0MUT_, which is not activated by 2-AG) into the aPVT and implanted a fiberoptic above the medial NAc shell (Fig 3N). We detected a progressive increase in the reward engagement-induced GRAB_eCB2.0_ signal from day 1 to day 5, and no change in basal GRAB_eCB2.0MUT_ signal (Fig 3O-Q). Compared to GRAB_eCB2.0MUT_ controls, we detected a significant increase in GRAB_eCB2.0_ signal on day 5 of reward conditioning. Furthermore, These differences in fluorescence were not due to overt changes in engagement behavior between the two groups (Fig 3R). We then tested whether 2-AG binds to aPVT terminals following exposure to aversive stimuli in a Pavlovian fear conditioning/extinction paradigm. Similar to our results gather with local NAc expression of GRAB_eCB2.0_ (Fig 1), foot shock elicited a robust increase in GRAB_eCB2.0_ signal while the shock predictive cue elicited no change in GRAB_eCB2.0_ signal during fear recall (Fig S10A-G).

Lastly, we sought to identify an in vivo correlate to the DSE phenomenon (neural depolarization induced production of eCBs) demonstrated in Figure 1. Specifically, we examined whether we could evoke 2-AG production and release through photo-stimulation of NAc neurons in vivo. Our electrophysiological results suggested that most of the phasic 2-AG release in the NAc is mediated via D_2_R+ neurons, which have been shown to co-express the opioid peptide Enkephalin with 95% overlap with D_2_R expression ^7,44^. Thus, we injected Pro-Enkephalin Cre (Penk-Cre) mice with the red-shifted channelrhodopsin AAV5-DIO-ChRimson-tdTomato in the medial NAc shell along with an implanted fiber optic, and virally expressed GRAB_eCB2.0_ in the aPVT (Fig 3S). 20hz 10s photo-activation of NAc Penk neurons were sufficient to elicit a significant increase in GRAB_eCB2.0_ signal, indicating in vivo mobilization and release of 2-AG and binding to aPVT terminals (Fig 3T). Overall, these findings demonstrate that in vivo release of 2-AG in the NAc functionally tunes engagement in reward-seeking behavior through time-locked inhibition of aPVT terminals.

### Encoding of behavioral engagement by aPVT-NAc^Penk^ neurons

Thus far, our evidence supports the role of 2-AG-CB_1_R signaling in filtering excitatory aPVT^NTS^input to the NAc to control reward-seeking, and that the NAc D_2_R/Penk neurons produce 2-AG. Given that we have demonstrated that reward-seeking-induced 2-AG release occurs in an activity dependent manner, we next asked how engagement in reward-seeking is encoded by the activity discrete ensembles of NAc Penk neurons. Mice were administered Pavlovian reward conditioning in a custom-built linear track behavioral corridor which allowed for multi-plane video recordings of animal behavior. To garner unbiased quantifiable parameters and features of engagement in reward seeking behaviors, we used a custom SLEAP pose estimation algorithm to identify and track specific locations on the mouse’s body (i.e., points of interest) during reward conditioning (Fig 4A) ^45^. A supervised machine learning algorithm was subsequently employed to identify frame-by-frame instances of sipper port engagement, sipper port disengagement, walking, rearing, and grooming (Fig 4B-H).

**FIGURE 4.**
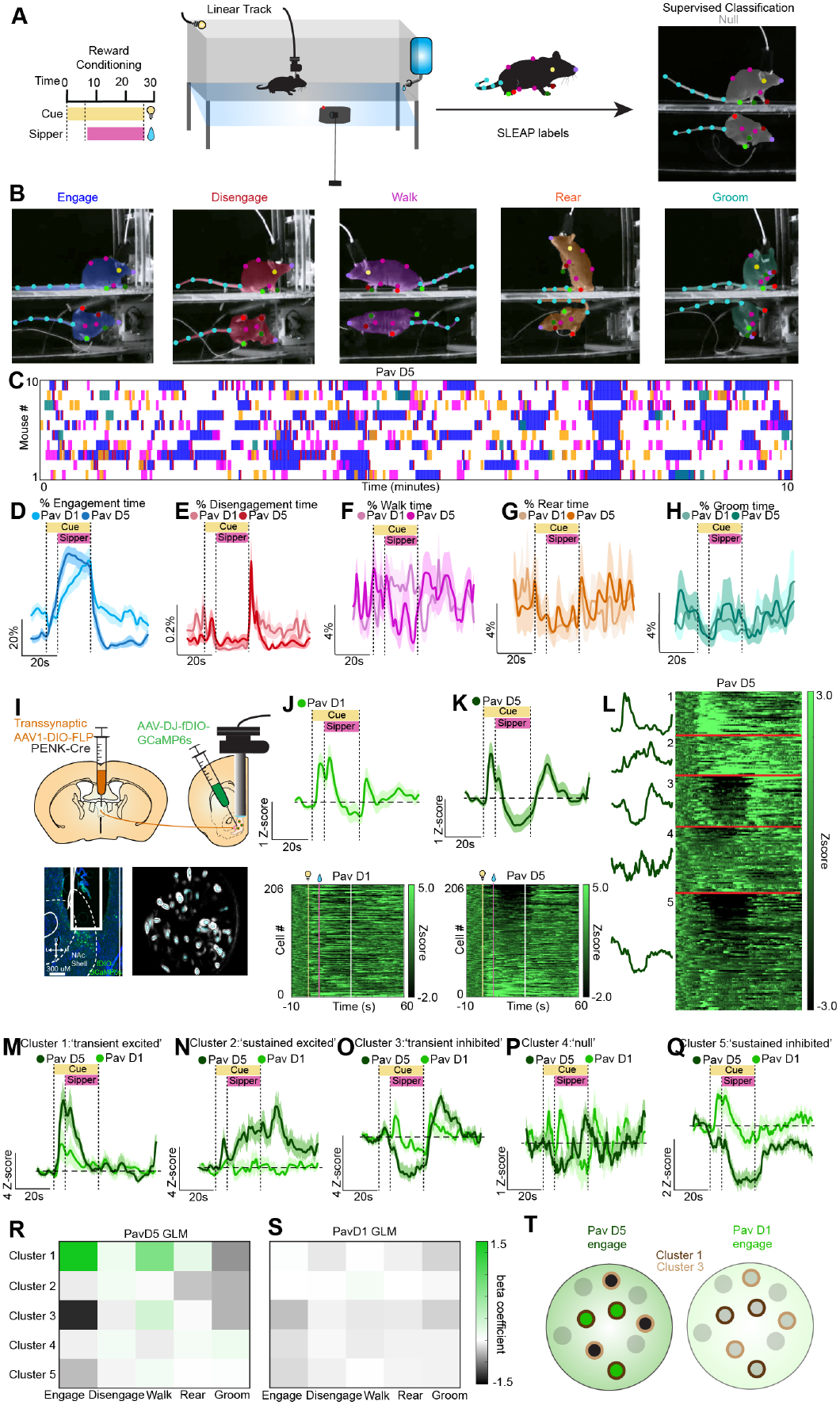
Encoding of behavioral engagement by aPVT-NAc^Penk^ neurons. Schematic for Pavlovian reward conditioning in the Linear Track and subsequent SLEAP labeling and supervised classification of behavior (A). Example frames of 5 classified behaviors: sipper port engagement, disengagement, walk, rear, and groom (B). Ethogram of the five classified behaviors from each of the 10 mice over a 10 minute epoch on day 5 of Pavlovian reward conditioning (C). Cue-aligned traces of % time engaging with sipper port (D), disengaging from sipper port (E), walking (F), rearing (G), and grooming (H) on PavD1 and PavD5. Schematic for trans-synaptic capture of NAc Penk neurons that receive aPVT input for subsequent 1-photon recordings of GCaMP6s of aPVT-NAc^Penk^ neurons and GRIN lens placement and representative maximum-projection image aPVT-NAc^Penk^ neurons (I). Average cue-aligned trace and heatmaps for 206 matched neurons from day 1 (J) and day 5 (K) of Pavlovian reward conditioning (N=10). Hierarchical clustering revealed 5 distinct clusters of aPVT-NAc^Penk^ neurons based on cue-aligned calcium signal on PavD5 (L). Traces of 5 unique cue-aligned clusters of aPVT-NAc^Penk^ neurons on day 5 of conditioning, was well as their neuron-matched traces from day 1 of conditioning (M-Q). Heat plot showing beta coefficients of a Generalized Linear Model trained on each PavD5 behavior throughout the 70 second window and fit to the activity of each cluster during the corresponding time window (R). Same as in (S) but for PavD1 behaviors (P). Graphic summarizing results of the GLM, depicting positive encoding of engagement in reward-seeking behaviors by aPVT-NAc^Penk^ cluster 2 and negative encoding by aPVT-NAc^Penk^ cluster 3 (T).

To understand how these behaviors were encoded at the neural level, we employed a strategy to record the activity of NAc D_2_R+/Penk neurons that specifically receive input from the aPVT (NAcaPVT). We leveraged a recently developed AAV1 virus (AAV-trans) that shows strong anterograde transsynaptic infectivity to drive expression of GCaMP in Penk neurons that receive aPVT input ^46,47^. Following the characterization of its labeling (Fig S11A,B), we co-injected Penk-Cre mice with AAV-trans-DIO-FLP into the aPVT combined with AAV-DJ-fDIO-GCaMP6s into the medial NAc shell to induce expression of GCaMP only in NAc Penk neurons that specifically receive aPVT afferent input. We then implanted a 0.6 mm GRIN lens over the medial NAc shell (Fig 4I). Using the spatial footprints of the neurons in the field of view, we matched 206 aPVT-NAc^Penk^ neurons between day 1 and day 5 of Pavlovian reward conditioning. Matched neurons from day 1 and day 5 of conditioning displayed remarkably similar dynamics (e.g., both cue- and sipper-delivery aligned excitation) as indicated when monitoring the summed Z-scored activity of all identified neurons. However, this averaged signal masked remarkable heterogeneity of individual neural responses both within each day, and between tracked cells across days (Fig 4J,K).

To gain a deeper understanding of this neural functional heterogeneity, we used PCA-based dimensionality reduction followed by hierarchical clustering to partition the neurons into functional clusters ^48,39^. These analyses were done on the Z-scored fluorescence activity of the cue-aligned calcium trace on day 5 of conditioning, and neurons were subsequently matched to their day 1 profiles. Using these approaches, we found that the neurons were partitioned into 1 of 5 total clusters (Fig 4L-Q, FigS11B-C, see methods). Cue cluster 1 displayed strong but transient activation to both cue and sipper delivery (‘transient excited’) (Fig 4M). Cue cluster 2 had a slower rise time in activity but more sustained increased dynamics lasting the duration of the duration of the 60 sec time window following cue delivery (‘sustained excited’) (Fig 4N). Cue cluster 3 showed both negative and positive deflections that closely mirrored the activity of aPVT terminals, suggesting the population could be entrained to the activity of these projections (‘transient inhibited’) (Fig 4O). Cue cluster 4 is principally a null cluster, as the average Z-score never peaks about 1 or dips below -1 (‘null’) (Fig 4P). Cue Cluster 5 is the primary inhibited cluster, showing decreased activity during both cue delivery and the subsequent reward (‘sustained inhibited’)(Fig 4Q). To substantiate cluster identities, a Support Vector Machine (SVM) was trained on PavD5 PC coordinates and cluster labels, which demonstrated accurate decoding of cluster identity for unshuffled day 5 data but not for day 5 shuffled or day 1 shuffled/unshuffled (Fig S11D). Notably, in a reward/threat cue-discrimination task, there was little overlap between ensembles that responded to rewarding vs threatening cues, suggesting that unique ensembles are involved in encoding engagement in reward-seeking and defensive freezing behaviors (Fig S13).

Simultaneous capture of 1p-recordings of aPVT-NAc^Penk^ GCaMP6s activity and high resolution Linear Track videos for SLEAP classification enabled us to determine how engagement in reward-seeking behaviors is encoded by changes in the activity of this neural population. Thus, we employed a Generalized Linear Model (GLM) fit of fluorescence activities of the 5 clusters, trained on frame-by-frame instances of the 5 aforementioned behaviors throughout the same 70 second window used for analysis of our 1-photon imaging data (10 seconds before to 60 seconds after the cue)(Fig S12). Of all behaviors across both days of Pavlovian conditioning, we identified 2 beta coefficients with an absolute value >1, belonging to the fit of PavD5 engagement to cue cluster 1 (transient excited) (β=1.483) and cue cluster 3 (transient inhibited) (β=-1.2476) (Fig 4R,S, Fig S13A-J). In this case, these coefficients represent the change in value of the response variable (cluster activity) given a 1 unit change in the predictor (behavior). Taken together, these results indicate that engagement in reward-seeking behaviors is encoded by the activity of specific aPVT-NAc^Penk^ ensembles (Fig 4T).

### Discrete ensembles of NAc_aPVT_ Penk neurons are entrained to aPVT terminal activity and endogenous cannabinoid release

Qualitatively, we saw that the activity of specific clusters of aPVT-NAc^Penk^ neurons resembled either aPVT^NTS^-NAc terminal activity or aPVT terminal GRAB_eCB2.0_ signal. To assess whether these signals were entrained to each other, we time-aligned the signals and ran a Hilbert analysis to decompose constituent waveforms and calculate instantaneous phase. These data were subsequently analyzed with a Rayleigh test for circular uniformity (Fig S14 A-J) ^49,50,51^. This test assesses the phase offset between time varying signals, and has been used to correlate single unit neural activity to local field potentials (Fig 5A) ^52^. When the clusters were aligned to aPVT terminal activity, we found the highest degree of entrainment (R=0.9353) between the ‘transient inhibited’ cluster (3) and the aPVT terminals with a phase offset of -21.11 degrees, suggesting the activity of the cluster slightly lags behind the terminal activity (Fig 5B,C). When the cluster activity was aligned with the GRAB_eCB2.0_ signal, we found the greatest degree of entrainment with the ‘sustained excited’ cluster (2) (R=0.8922) with a phase offset of -1.14 degrees, suggesting that the signals emerge contemporaneously Fig 5D,E). Overall, these data suggest that specific ensembles of aPVT-NAc^Penk^ neurons are entrained to aPVT^NTS^-NAc terminal activity or aPVT terminal GRAB_eCB2.0_ signal.

**FIGURE 5.**
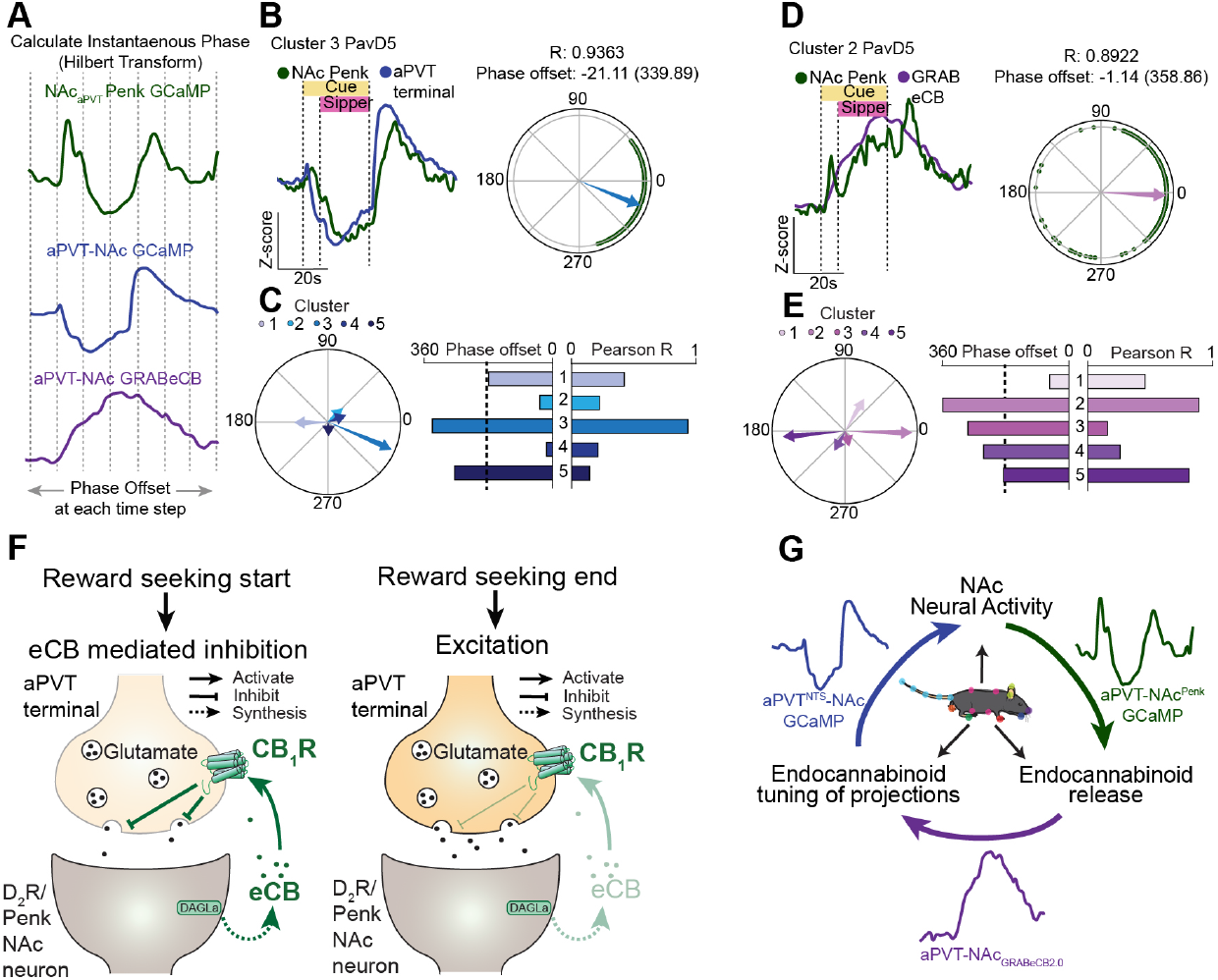
Discrete ensembles of aPVT-NAc^Penk^ neurons are entrained to aPVT terminal activity and endogenous cannabinoid release. Schematic depicting alignment of aPVT-NAc^Penk^ GCaMP, aPVT^NTS^-NAc GCaMP, and aPVT-NAc GRAB_eCB2.0_ signals for Hilbert analysis and Rayleigh plots. Overlay of aPVT-NAc^Penk^ cluster 3 GCaMP6s signal with aPVT^NTS^-NAc terminal GCaMP6s signal from PavD5 and corresponding Rayleigh plot for circular uniformity (R=0.9363) (B). Summary plot of Hilbert and Rayleigh analyses depicting Pearson R and phase offset values for each cluster’s entrainment to aPVT^NTS^-NAc GCaMP signal (C). Overlay of aPVT-NAc^Penk^ cluster 2 GCaMP6s signal with aPVT-NAc GRAB_eCB2.0_ signal from PavD5 and corresponding Rayleigh plot for circular uniformity (R=0.8922) (D). Summary plot of Hilbert and Rayleigh analyses depicting Pearson R and phase offset values for each cluster’s entrainment to aPVT-NAc GRAB_eCB2.0_ signal (E). Summary schematic depicting reward-seeking induced eCB production by NAc Penk neurons and subsequent retrograde action at aPVT terminals, and excitation of aPVT terminals upon disengagement (F). Summary schematic depicting the reciprocal relationship between aPVT-NAc^Penk^ neuron activity, eCB release and presynaptic binding, eCB tuning of excitatory input from the aPVT, and entrainment of aPVT-NAc^Penk^ neurons to aPVT terminal activity (G).

## 3 DISCUSSION

The behavioral role of eCB signaling on distinct neural circuits *in vivo* has been difficult to probe due to widespread expression of this signaling system and a lack of tools to assess eCB dynamics. Using a suite of electrophysiology, fiber photometric, optogenetic, CRISPR/Cas9 deletion, and 1-photon imaging approaches, we uncovered a novel eCB mechanism for regulation of reward-seeking behaviors through modulation of a genetically and anatomically defined thalamo-striatal circuit (Fig 5F). While multiple excitatory projections to the NAc expressed the CB_1_R receptor, we found the highest density was localized in aPVT projectors which express the neuropeptide neurotensin. Electrophysiological characterization of this circuit revealed that these inputs are regulated by both tonic and phasic 2-AG produced by D_2_R/Penk neurons. *in vivo*, the activity of these projections was negatively correlated with engagement in both reward seeking behaviors as well as conditioning freezing. However CRISPR/Cas deletion of CB_1_R from aPVT NTS neurons was only sufficient to attenuate reward-seeking behaviors (and concomitant aPVT terminal inhibition), leaving conditioned freezing behaviors intact. Given that activity dependent production of 2-AG was primarily evoked from D_2_R/Penk neurons, we used a transsynaptic 1-photon imaging strategy to demonstrate that the activity of specific ensembles of clusters of these aPVT-NAc^Penk^ neurons encodes engagement in reward-seeking behaviors, and that specific clusters are entrained to either aPVT terminal activity or 2-AG binding to aPVT terminals.

Multiple *in vivo* electrophysiology studies have demonstrated that reward seeking/consumption elicits an increase in the firing rates of PVT neurons ^53,54,55^. This is at odds with the results of several fiber photometry studies demonstrating inhibition of PVT terminals in the NAc during epochs of reward-seeking. This suggest that local modulation of PVT terminal activity within the NAc must be responsible for the resultant inhibition of these projections. Our data identify eCBs in the NAc as cruical regulators of reward-seeking behaviors through inhibition of excitatory input from the aPVT.

Recent reports investigating functional discrepancies between the aPVT and pPVT have suggested that aPVT projections to the NAc encode the termination of goal pursuits ^32,33^. Our framing of the activity of these projections encoding behavioral engagement through inhibition and disengagement through excitation are largely consistent with this finding, although our studies expand the scope of this model to include engagement in aversive behaviors (such as conditioned freezing) as well. Our finding that deletion of the CB_1_R from aPVT neurons was sufficient to attenuate reward engagement-induced inhibition but not freezing-induced inhibition of aPVT-NAc terminals was surprising. This suggests that while inhibition of aPVT-NAc terminals may be a conserved feature of behavioral engagement, the mechanism driving this inhibition could be dependent on either stimulus valence or a property of the specific behavioral output.

Our electrophysiological data demonstrated that depolarization-induced suppression of excitation (DSE) is principally expressed at aPVT^NTS^-NAc Penk synapses. A recent report for the first time demonstrated an *in vivo* correlate of this phenomenon in the CA1 area of the hippocampus using GRAB_eCB2.0_ ^56^. Using this tool in combination with the red-shifted channelrhodopsin ChRimson, we were able to evoke an increase in GRAB_eCB2.0_ signal following NAc Penk neuron activation, ostensibly evoking a DSE-like phenomenon. Further studies using 2-photon imaging will be needed to conclusively demonstrate *in vivo* DSE at these synapses.

While prior work has demonstrated the feasibility of 1- and 2-photon recordings of NAc neurons in a genetically specific manner, no studies to date have recorded from NAc neurons with respect to both afferent connectivity and genetic identity ^57,58,59^. Our transsynaptic approach allowed us to specifically image from NAc Penk neurons which receive direct monosynaptic input from the aPVT, allowing us unprecedented granularity, which is uniquely important in the NAc given the well characterized functional diversity of NAc neurons ^7^. This approach combined with dimensionality reduction and hierarchical clustering yielded 5 unique ensembles of neurons based on their reward cue-aligned responses. While some clusters (e.g. 1) had clear cue/reward evoked inhibition / excitation, many of the response patterns were complex and evolved across the 60 sec following cue presentation. By utilizing novel pose estimation and machine learning algorithms to garner quantitative metrics of behavioral states and transitions, we demonstrated that the activity of specific clusters of aPVT-NAc^Penk^ neurons encodes engagement in reward seeking behaviors. Furthermore, using signal processing analyses (Hilbert transform and Rayleigh test), we demonstrated that unique clusters of aPVT-NAc^Penk^ neurons are entrained either to NAc confirmed that these clusters were significantly entrained to these respective signals. Of note, a caveat to these approaches is that such analysis was performed between mice rather than within mice. A more complex combination of 2-photon imaging, spatial light modulation, and terminal imaging within a single animal would be needed to conclusively link these datasets.

Overall this report uncovers a fundamental molecular mechanism that involves localized NAc 2-AG production in the regulation of engagement in reward-seeking behaviors through neuromodulatory gain-control of an anatomically and genetically defined thalamo-striatal circuit (Fig 5H,I). These data demonstrate a causal link between single cell neural activity (aPVT-NAc^Penk^), neuromodulator release (GRAB_eCB2.0_), neuromodulatory tuning of projections (aPVT^NTS^-NAc GCaMP), and ultimately how each of these neural mechanisms regulate behavioral engagement during reward seeking. (Fig 5G). These results provide a novel framework for understanding the basic neurobiology of reward-seeking, while also establishing how neuromodulatory signaling influences unique endophenotypes implicated in the pathophysiology of substance use disorders.

## 4 MATERIALS AND METHODS

### Animals

Adult (16-35g, 12-52 weeks) male and female WT, NTS-Cre, PENK-Cre, and Ai14 mice were group housed, given adlibitum access to food and water, and maintained on a 12 hour light dark cycle (lights off at 9:00 AM, lights on and 9:00 PM). Mice were bred in a barrier facility and transferred to the holding facility at least 1 week before the start of surgical interventions or experiments. Littermate-matched controls were used in manipulation studies (optogenetics, CRISPR, behavioral pharmacology). All animals were drug and test naïve, unless otherwise noted. For all reward assays, mice were water restricted to 90% of baseline body weight for at least 5 days before behavioral testing. If body weight dipped below 85% of baseline weight, mice were removed from water restriction. Statistical comparisons did not detect any sex differences, and therefore male and female mice were combined for all of the data presented in this manuscript. All animals were monitored daily by either facility staff or experimenter throughout the duration of the study. All procedures were approved by the Animal Care and Use Committee at the University of Washington, and conformed to US National Institutes of Health guidelines.

### Drugs

For all behavior experiments, drugs were injected I.P. and prepared in a 17:1:1:1 vehicle consisting of saline (17), ethanol (1), kolliphor (1), and DMSO (1). Drugs were administered at the following concentrations: SR141716A (Sigma): 10mg/kg, DO34 (Sigma): 20mg/kg, JZL184 (Sigma): 5mg/kg. SR141716A and JZL184 were injected 90 minutes before behavioral testing, and DO34 was injected 120 minutes before behavioral testing.

### Stereotaxic Surgery

Mice were anaesthetized in an induction chamber (1%-4% isoflurane) and placed into a stereotaxic frame (Kopf Instruments, model 1900) where they were mainlined at 1%-2% isoflurane. For all viral injections, a Hamilton Neuros Syringe was used, and virus was infused at a rate of 50-100 nl/minute. For retrograde tracing studies, 150nL of AAV2-retro-DIO Cre was injected into the medial NAc shell of Ai14 mice at two D/V coordinates per hemisphere, A/P 1.7, D/V 4.4 and 4.1, M/L 0.65. For fiber photometry experiments, AAV5-DIO-GCaMP6s (Addgene) or AAV1-hSyn GRABeCB was injected into the aPVT of NTS-Cre or WT mice respectively at a 10 degree angle, with final coordinates of A/P -0.4, D/V -3.75, M/L 0. For CRISPR/ photometry experiments, AAV5-DIO-GCaMP6s was mixed in a 2:1 ratio with AAV1-FLEX-sgCNR1-SaCas9 or AAV1-FLEX-sgROSA26-SaCas9 and then injected into the aPVT of NTS-Cre mice. Three weeks after viral injection, a 400μM fiber optic with a stainless steel ferule (Doric) was implanted in the medial NAc shell at A/P +1.7, D/V -4.2, M/L +/-0.65. All implants were secured using Metabond (Parkell). For optogenetics experiments, 500nl of AAV5-DIO-ChR2-eYFP or AAV5-DIO-PPO-eYFP (Addgene) was injected into the aPVT of NTS-Cre mice. Three weeks after injection, bilateral 200μM fiberoptics were implanted over the medial NAc shell at a 10 degree angle (final coordinates of A/P +1.7, D/V -4.2, M/L +/-0.65). For 1-photon imaging experiments, 500nL of AAV1-trans-DIO-FLP was injected into the aPVT of PENK-Cre mice. Three weeks later, 250 nL of AAV5-DIO-GCaMP6s was injected in the medial NAc shell at two D/V coordinates per hemisphere, A/P 1.7, D/V 4.4 and 4.1, M/L 0.65. During the same surgery, a 600μM cuffed GRIN lens (Inscopix) was implanted at coordinates of A/P 1.7, D/V 4.15 M/L 0.65. Three weeks after viral injection and lens implantation, we attached a baseplate to an Inscopix miniscope to identify the optimal imaging plane. The baseplate was subsequently attached to the implant with Metabond. All mice were given at least 3 weeks of recovery time post-surgery before experimentation.

### Tissue Processing

For all *in vivo* optogenetics, fiber photometry, or 1-photon imaging experiments, mice were transcardially perfused with a 4% paraformaldehyde solution in phosphate buffered saline. Heads were subsequently removed and immediately placed in to conical tubes containing the same solution and allowed to set for 24-72 hours. After this time, fiberoptic implants were removed along with the remainder of the skull, and brain were put into a cryoprotectant 30% w/v sucrose solution in PBS. Once brains had sunken to the bottom of the tubes (typically around 72 hours), brains were mounted on cutting blocks using O.C.T. (Sakura Finetek) and sliced in 40μm sections on a Leica CM1900 cryostat at - 18 degrees C. Slices were then placed into PBS and either immediately mounted on SuperFrost Plus slides (Fisher) or stored for up to 3 days. Mice with misplaced fiber optic implants were excluded from the study.

### In situ hybridization using RNAscope

For quantification of mRNA transcripts using RNAscope (ACD Bio), mice were briefly anesthetized using isoflurane and subsequently rapidly decapitated. Brain were immediately removed from the skull and were placed into a beaker containing 2-methylbutne that had been sitting on dry ice for >20 minutes. After 30-60 seconds of submerging, brains were removed and placed into conical tubes in a -80 degree C freezer for long term storage. Prior to sectioning, brains were placed in the cryostat compartment for >30 minutes to allow them to come up to temperature. Following mounting using O.C.T., brains were slice at 16μm and directly mounted onto SuperFrost Plus slides. Slides were stored at -80 degree C.

FISH was performed according to the RNAscope 2.0 Fluorescent Multiple Kit User Manual for Fresh Frozen Tissue (Advanced Cell Diagnostics) as previously described. Briefly, slides were fixed in 40% PFA and then serially dehydrated using increasing concentrations of ethanol (50%, 75%, 100% and 100%). Slices were then treated with Protease IV and allowed to incubate in a 40 degree C hybridization oven for 30 minutes. Following serial rinses in a wash buffer, slices were incubated with probes directed against Cre (312281-C3), CB1 (457341), or NTS for two hours (420441-C2). Samples subsequently went through sequential amplification steps, and lastly application of Opal dyes corresponding to each channel (TSA Vivid Fluorophore 520 (PN 323271), TSA Vivid Fluorophore 570 (PN 323272) and TSA Vivid Fluorophore 650 (PN 323273)). Slides were counterstained with DAPI, and coverslips were mounted with Vectashield Hard Set mounting medium (Vector Laboratories).

### Image acquisition and analysis

For histological verification of fiberoptic/lens placement, images were obtained on a Leica DM6B epifluorescent microscope at 10x or 20x magnification. For RNAscope analysis, images were collected on an Olympus Fluoview 3000 at 20x or 40x magnification. Z-stacked images were taken with optical section thickness of 2-3μM. Images were subsequently analyzed using either HALO or QuPath software. For either analysis, brain region boundaries were drawn in accordance with the Allen Brain Institute mouse reference atlas. For HALO, DAPI stained nuclei were used to mark cells, and cell boundaries were set to 8 μM radius from nucleus center. Thresholds were set for detection in each channel and this threshold was applied to all analyzed images in a given cohort. In order to account for background noise, a neuron had to have >2 transcripts in order to be classified as positive for a specific marker. For QuPath analysis, DAPI was similarly used to define neurons and cell boundaries were set to 8 μM radius from nucleus center. Individual channels were set via the classifier module, and these classifier setting were held consistent for all images within a cohort. The classifiers were set by individually sorting through each cell automatically registered by the QuPath software and identifying the first ‘false positive’ neuron, using this as the threshold cut off for subsequent identification of positive neurons.

### Electrophysiology

Coronal brain slices were prepared at 250 μM on a vibrating Leica VT1000S microtome using standard procedures. Mice were anesthetized with Euthasol (Virbac, Westlake Texas), and transcardially perfused with ice-cold and oxygenated cutting solution consisting of (in mM): 93 N-Methyl-D-glucamine (NMDG), 2.5 KCL, 20 HEPES, 10 MgSO4-7H20, 1.2 NaH2PO4, 0.5 CaCl2⍰H20, 25 glucose, 3 Na+-pyruvate, 5 Na+-ascorbate, and 5 N-acetylcysteine. Following collection of coronal sections, the brain slices were transferred to a 34°C chamber containing oxygenated cutting solution for a 10-minute recovery period. Slices were then transferred to a holding chamber consisting of (in mM) 92 NaCl, 2.5 KCl, 20 HEPES, 2 MgSO4⍰7H20, NaH2PO4, 30NaHCO3, 2 CaCl2-2H20, 25 glucose, 3 Na-pyruvate, 5 Na-ascorbate, 5 N-acetylcysteine and were allowed to recover for ≥ 30 min. For recording, slices were perfused with oxygenated artificial cerebrospinal fluid (ACSF; 31-33°C; 300-303 milliosmols) consisting of (in mM): 113 NaCl, 2.5 KCl, 1.2 MgSO4⍰7H20, 2.5 CaCl2-6H20, 1 NaH2PO4, 26 NaHCO3, 20 glucose, 3 Na+-pyruvate, 1 Na+-ascorbate, at a flow rate of 2-3ml/min. Stocks of 50mM SR1 or 5mM JZL184 were made in DMSO and diluted 1:5000 in ACSF or HEPES for a final concentration of 10μM and 1μM, respectively. 0.05% w/v Bovine Serum Albumin (Sigma-Aldrich) was added to ACSF or HEPES to help the drugs from precipitating out of solution. For all drug experiments, slices were incubated in a HEPES bath containing the drugs for >30 minutes before transferring over to the slice rigs for recording.

NAc neurons were initially voltage clamped in whole-cell configuration using borosilicate glass pipettes (2-4MΩ). filled with internal solution containing (in mM): 125 K+-gluconate, 4 NaCl, 10 HEPES, 4 MgATP, 0.3 Na-GTP, and 10 Na-phosphocreatine (pH 7.30-7.35). The patch pipette also included 50 μM picrotoxin to block GABAA currents. D_1_R+ medium spiny neurons (MSNs) were identified by tdTomato fluorescence. For all D_1_R-neurons, multiple electrophysiological metrics were recorded to ensure we were specifically recording from MSNs. This included capacitance, membrane resistance, resting membrane potential, sag current, afterhyperpolarization current, and evoked action potential firing. Multiple publications have identified that these features differ between MSNs and interneurons, and any neurons that had electrophysiological features of interneurons were excluded from the dataset (Figure S1) ^60,61^. Sag current was recorded following a 1 second 100pA hyperpolarizing current, and the amplitude of the deflection below the stable baseline was calculated. For afterhyperpolarization current, we analyzed the amplitude of the inward current immediately following a train of action potentials induced by a 500pA current step. Action potential firing was monitored during 40 consecutive successive sweeps of currents starting at -100pA and increasing by 20pA each time. Following break-in to the cell, we waited ≥ 3 minutes to allow for exchange of internal solution and stabilization of membrane properties. Neurons with an access resistance of > 30MΩ or that exhibited greater than a 20% change in access resistance during the recording were not included in our datasets.

### *Ex vivo* optogenetics

For electrophysiological interrogation of the aPVTNTS-NAc circuit, mice were injected with 500 μL of AAV5-DIO-ChR2(H134R)-eYFP into the aPVT. 3-5 weeks of viral expression was allowed prior to sacrificing the mice. For all experiments using SR141716A or JZL184, corresponding vehicle recordings were obtained from the same mice. For optogenetic recordings of input/output curves, we used a Thorlabs LEDD1B T-Cube driver and obtained separate recordings of 470nm wavelength oEPSCs at 7 output levels corresponding to 5.7, 2.7, 1.6, 1, 0.5, 0.2, and 0.1 mW of LED intensity. PPR recordings of oEPSCs were obtained in voltage-clamp with an inter-stimulus interval of 50ms. PPR is reported as a ratio between the amplitude of the second oEPSC divided by the first. Recordings of DSE were obtained following at 10 s voltage step to +30mV. A baseline of 10 oEPSCs were taken prior to the depolarizing step, and all data is plotted as an oEPSC amplitude normalized to the baseline period. A light exposure time of 2ms was used for all optogenetic experiments.

### Fiber photometry

Fiber photometry recordings were obtained throughout the duration of all behavioral tests presented in this manuscript. Prior to recordings, a fiberoptic cable was attached to the fiberoptic implant using a plastic or ceramic ferrule sleeve (Doric, ZR2.5). For all photometry recordings, we used a Tucker-Davis Technologies RZ10x processor. A 531-Hz sinusoidal 470 nm LED light (Lx465) was used to excite GCaMP6s and evoke Ca2+-dependent emission. A 211-Hz sinusoidal 405 LED (Lx405) light was used as the Ca2+-independent isosbestic control emission. For GRAB_eCB2.0_ recordings, we used a 490 nm LED for excitation and a 435 nm LED as an isosbestic control, as prior studies indicated that the peak excitation wavelength for this sensor is slightly red-shifted ^31^. LED intensity was measured at the tip of the cable with a dummy implant attached and adjusted to 30μW before each day of recording. GCaMP6s fluorescence traveled through the same optic fiber before being bandpass filtered (525 ± 25 nm, Doric, FMC4), transduced by a femtowatt silicon photoreceiver (Newport, 2151) and recorded by a real-time processor (TDT, RZ10). The envelopes of the 531-Hz and 211-Hz signals were extracted in real-time by the TDT program Synapse at a sampling rate of 1017.25 Hz. For ChrimsonR stimulation experiments, a 635nm laser was used with a custom filter cube (Doric) at 5-7mW of intensity to deliver 20hz pulsed light through the same fiberoptic cable used for photometry recordings.

### Photometry Analysis

Custom MATLAB scripts were developed for analyzing fiber photometry data in context of mouse behavior and can be accessed via GitHub. The isosbestic 405/435nm excitation control signal was scaled to the 470/490nm excitation signal, then this refitted 405/435nm signal was subtracted from the 470/490 nm excitation signal to remove movement artifacts from intracellular Ca2+-dependent GCaMP6s/GRAB_eCB2.0_ fluorescence. Baseline drift was evident in the signal due to slow photo-bleaching artifacts, particularly during the first several minutes of each hour-long recording session. A double exponential curve was fit to the raw trace and subtracted to correct for baseline drift. After baseline correction, Δf/f was calculated by subtracting the median baseline subtracted signal from the baseline subtracted signal and dividing by the median raw signal. All photometry data presented in this manuscript is Z-scored to a 10-20 second window prior to cue onset.

### Behavioral apparatuses and analysis

#### Pavlovian reward conditioning

Mice were habituated to a Med Associates operant conditioning box (MED-307W-D_2_R) for 3 days (30 minutes per session), during which a sipper containing a 10% sucrose solution was extended into the chamber. Mice were subsequently trained in Pavlovian Reward Conditioning, in which the houselight was illuminated 6 seconds prior to sipper extension, which lasted for 20 seconds, followed by sipper retraction and houselight shut-off. Licks were recorded via a contact lickometer and registered in MedPC. The cue/sipper pairing was presented in 20 pseudorandom trials with ITIs of 30, 60, 90, and 120 seconds. For all photometry experiments, a 1080P webcam was mounted above the chamber for subsequent mouse behavior analysis. For engagement in reward-seeking behaviors, we analyzed the % of time the mouse spent pawing, sniffing, licking, or poking at the reward port in 3 defined time windows: 0-6 seconds (cue period, pre-sipper), 0-26 seconds (entire cue period), or 26-46 seconds (post-cue/sipper time period). These behaviors were analyzed in a blinded fashion. The same Pavlovian protocol (identical cue/reward pairings, ITI, sucrose concentration) was used for conditioning in the Linear Track (see below).

#### Operant conditioning

Using the same apparatus for Pavlovian reward conditioning, we trained mice on FR1, FR3 and PR schedules of reinforcement. Animals nosepoked into illuminated nose-poke ports, randomly designated as ‘active’ or ‘inactive’. Following a second delay, an active nosepoke resulted in cue light presentation and subsequent protraction of the sipper, which stayed protracted for 20s. In a FR1 schedule of reinforcement, one nosepoke elicited sipper protraction. Following 3 days of FR1 training, mice were trained in a FR3 schedule which necessitated 3 consecutive active pokes to elicit sipper protraction. Finally, in the PR task, the number of nosepokes required to elicit sipper protraction escalated exponentially as previously described ^62^.

#### Pavlovian fear conditioning and extinction

In separate Med Associates chambers (NIR-022MD), mice were trained in Pavlovian fear conditioning. The chamber consisted of a conductive grid floor (29.53 × 23.5 cm) illuminated by an IR light. Mice were conditioned to 10 20khz tones lasting 10 seconds, co-terminating with a 2 second .5mA shock. Tones were presented in a pseudorandom pattern. For fear extinction, mice were exposed to 20 20hz tones lasting 20 seconds for five consecutive days. Freezing behaviors was calculated manually by an experimenter blinded to animal group and genotype.

### *in vivo* optogenetics

For all experiments using ChR2 we used a 20hz stimulation frequency with a power output between 6 and 8 mW (measured at fiber tip). For all experiments using PPO we used a 10hz stimulation frequency with a power output between 8 and 10 mW.

#### Pavlovian reward conditioning

For reward conditioning experiments, mice were exposed to the identical training regimen as described above. For modulation of reward-seeking behaviors, we used two separate stimulations paradigms for ChR2. On counter-balanced days, mice either received 20hz stimulation throughout the duration of the cue period (26s) or received closed-loop 20hz stimulation in which each lick triggered (via TTL) a 2 second stimulation pulse. For PPO, we drove 10hz inhibition via a TTL pulse which began the stim pattern 5 seconds before house-light shut-off and continued for 20 seconds after houselight shut-off.

#### Pavlovian fear conditioning

For reward conditioning experiments, mice were exposed to the identical training regimen as described above. Mice received optical stimulation or inhibition on day 1 of extinction (fear recall day). For ChR2, mice received 20hz stimulation time-locked to cue presentation (20s). For PPO, mice received 10hz inhibition time-locked to cue presentation (20s).

#### Operant self-stimulation

Using the operant boxes described above, mice were trained in an FR1 schedule of reinforcement. One nosepoke either elicited a 2 second 20hz stimulation (ChR2) or 10hz inhibition (PPO) pulse.

### 1-Photon imaging

At least 1 week after baseplate attachment, mice were habituated to a Inscopix nVista 2.0 miniscope for 30-35 minutes for at least three days in the linear track, during which mice also had access to sipper containing a 10% sucrose solution. During the habituation phase, optimal field of view, focus, gain, and LED power were determined visually by assessing the activity of putative neurons in the field of view using the Inscopix data collection GUI. Gain values ranged between 2 and 6 and LED power was set between 20% and 50% of transmission range. Once determined, these parameters were held static for the remainder of the testing regimen.

Following this habituation period, mice were trained in Pavlovian reward conditioning as described above. Mice were scruffed and attached to the miniscope, and then immediately placed in the Linear Track. The Inscopix recordings were triggered directly by the MedPC program via a TTL pulse to ensure synchronization of event timestamps and calcium activity. The MedPC program was started 10s after starting the Linear Track camera, and videos were synced post-hoc via visible behavioral events (cue on, sipper protraction, lick, etc.). Animal rotations were compensated for by an automatic motorized commutator (Inscopix).

### Calcium imaging analysis

#### Initial Processing

All pre-processing was done using the Inscopix data processing software. Videos were initially spatially downsampled by 4x, followed by application of a bandpass filter (0.005 to 0.5 pixel-1). Lateral motion was corrected using the built in motion correction feature in the Inscopix data processing software. Motion corrected tiffs were subsequently exported for cell segmentation and further analysis. Using custom Matlab scripts (see GitHub), the motion corrected .TIFF video was then processed using the Constrained Non-negative Matrix Factorization approach (CNMFe), which has been optimized to isolate signals from individual putative neurons from microendoscopic imaging ^63^. Putative neurons were then manually sorted and registered using previously established criteria ^64^. For all analyses presented in this manuscript, the non-denoised, non-deconvoluted traces were used. For each cell, the raw fluorescence trace was Z-scored to a 10 second baseline period preceding the behavioral or task-related features presented in each figure.

#### Cell Matching

For matching of cells across days, we used the CellReg GUI as detailed in Sheintch et al 2017^65^. Briefly, maximum projection spatial footprints were extracted from the exported tiff files using a custom Matlab code. Cell registration occurred in 5 subsequent steps: 1). Spatial footprints from day 1 and day 5 of Pavlovian reward conditioning were loaded into the CellReg GUI 2). Field of view alignment was performed using rigid alignment, incorporating translations and rotations, with a maximal rotation of 30 degrees 3). Probabilistic modeling was run with a maximum distance of 12 microns 4). Initial cell registration was performed using spatial correlations 5). Final cell registration incorporated the probabilistic model from step 3 as well as the spatial correlations from step 4.

#### Principal component analysis and hierarchical clustering

For all Pavlovian reward conditioning imaging data presented in this manuscript, cells were sorted based on their day 5 activity and then back-matched to day 1 using custom matlab codes (Github). For Hierarchical clustering, a Principal Component Analysis was initially performed based on the activity of the neurons on Day 5 of Pavlovian reward conditioning over a one minute window coinciding with cue-onset for cue-associated activity and a ten second window for engagement-associated activity. Using built in Matlab Linkage function, an agglomerative hierarchical clustering algorithm was applied to the PCA of the day 5 activity. We set an a priori threshold that in order to be classified as a cluster, the number of neurons in said cluster must comprise >10% of the total population of recorded neurons (i.e. >20 of 206 neurons). As agglomerative hierarchical clustering starts with n clusters = n cells, we began by setting the cluster number equal to our a priori threshold (i.e. 10 clusters for 10% in each, assuming equal distribution among clusters). We then sequentially decreased cluster size by 1 until our a priori criteria were met.

### Cluster validation with a Support Vector Machine

SVM validation of cluster classification was performed in 5 sequential steps.

#### 1). Input data

PCA + hierarchical clustering (deterministic) was done using MATLAB on both day 5 and day 1 data, and then export PC coordinates and cluster labels. The cluster labels are based on the day 5 data and corresponding cell in day 1 gets the same cluster label. All data is exported as an excel file.

#### 2). Data preprocessing

The data and cluster labels are loaded and then split 80-20 into training and testing data, respectively. The 80% represent the training data and the remaining 20% is the testing data. The the data is then scaled by fitting sklearn’s standard scaler to the training data and then applying it to the testing data.

#### 3). Training the SVM

Used Sklearn’s SVC to perform multiclass classification using one versus one approach. Using 80% of the training data, we first performed cross validation using stratified k-fold and random search. The cross validation seem to suggest the linear kernel performed best across different numbers of PCs. We then used the identified hyperparameters.

#### 4). Classifying individual neuron activity

The fitted model is used to predict the classes for the testing data, as well as entirety of day 1 data. F1 score and accuracy are reported.

#### 5). Testing against pseudo-random data

The SVM was then fit to shuffled day 5 data (each time step values are shuffled), and then fit to day 5 data with shuffled cluster labels. In each case, NumPy’s permutation was used. Again, accuracy and f1 score was used to evaluate performance of the model.

### Phase analysis

The analysis requires at least two time series able to be loaded as NumPy arrays, they should have the same sampling rate and length. The analysis assumes each element correspond to each other. In our case, the time series was exported from MATLAB after preprocessing and linear interpolation to ensure sampling rate and length match. No further data preprocessing was necessary. The instantaneous phase of each time series is extracted using the Hilbert transform, creating an analytic signal, which is a complex-valued function that extends a real-valued function to the complex plane. The instantaneous phase is then obtained by taking the angle of this analytic signal. SciPy signal’s hilbert function in conjunction with NumPy was used to extract the phases. The phase difference between the two signals is computed by subtracting the instantaneous phase of one signal from the other. This difference is then wrapped to the range [-π, π] to ensure it falls within a single cycle.

To quantify the consistency of the phase difference across time, we use circular statistics. First we determined the mean resultant length (R) which represents the concentration of phase differences around the mean. It is calculated by 1). Converting each phase difference into a unit vector on a complex plane, 2). taking the mean of these vectors, and 3). calculating the magnitude of this mean vector. Second, we determined the mean phase difference (theta mean). This is the average angle of phase differences, which is calculated by taking the arctangent of the ratio of the mean sine to mean cosine of the phase differences. R values range from 0 to 1, where 0 indicates uniformly distributed phases and 1 indicated perfectly consistent phase differences. The analysis results were then plotted in a polar graph (Rayleigh plot), where each phase difference was plotted along with an arrow representing the R and mean theta values.

### Linear Track recordings

For high resolution point of interest and kinematic behavioral analysis, a linear track behavior corridor was utilized to capture dual perspectives of animal positions and behavior (bottom-up and side profile). The chamber was modified from the original published by Machado et al 2015 with a mirror positioned at a 45° angle below a clear Plexi-glass floor ^66^. A single high-frame rate camera was used to capture minute, quick animal actions during experimental windows (Basler ace camera: acA800-150um). The linear track was illuminated with 4 infrared (IR) floodlights and 2 supportive IR lights positioned around the chamber to brighten all angles of animal positions. All videos were recorded at 60 frames per second. The camera was operated via a custom python script which allowed for manipulation of session length, exposure, and capture rate (Github) outputting compiled mp4 videos for SLEAP analysis. The maximum session length was set to 40 minutes, although most recordings were stopped prior to this point, coinciding with the end of the behavioral assay.

### Machine learning behavioral tracking

Tracking and pose-estimation of videos within the linear track was performed using the SLEAP platform, an open-source computer vision system utilizing a custom convolutional neural network designed for variational animal behavior tracking ^45^. A custom, in-house tracking algorithm was developed for our linear track chamber using a subset of 1767 frames from 313 videos of adult (6 week to 6 month) 51% male and 49% female mice in different lighting conditions to ensure model robustness and versatility. This model was utilized to track 29 selected points of interest across all frames of all behavioral videos and were then exported as .h5 files for subsequent analysis. These 29 points were selected to represent the status of the animal from the bottom-up and side profiles in a single-animal model (e.g. nose-side profile and tail base-bottom profile). Of the total training frames, 50% were randomly selected by SLEAP and 50% were hand-picked for valuable frames to improve model accuracy. Final model precision and recall was 97.15% and 98.74%, respectively. An analysis package was used to export the positional data and calculate features for respective classification models.

### Supervised analysis of standard behaviors

Identification of core behavioral responses within the linear track was performed with an automated supervised ML algorithm to identify frame-by-frame instances of walking, rearing, grooming, and laying on belly. This model was trained on 1.4 million frames of naturalistic behavior within the linear track from a dataset of 45 videos consisting of adult (6 week to 6 month) 51% female and 49% male mice. Videos of animals were treated with either vehicle or some drug to expand the model versatility across different behavioral modifications. Model was trained on 1.35 million frames and separated test set revealed a final precision and recall of 98.3% and 97.7%, respectively. Further code was used to isolate behavioral event timestamps from the supervised model to align neural data when applicable.

### Engagement analysis

To evaluate mouse engagement with a sipper port, we built a pipeline that processes data from multiple experimental sessions. This pipeline takes three key inputs: (1) SLEAP joint output files in H5 format, containing joint coordinates of shape (2, 30, n) where n is the number of frames; (2) session names to be processed (e.g. PENKtrans-M2-D5); and (3) sipper port locations as pixel coordinates in an array of shape (2, 2). Our preprocessing stage involves three key steps. First, we detach metadata from the joint files to isolate pose data. Next, we extract the side head position, specifically the neck and nose coordinates, which are crucial for our analysis. Finally, we convert head and sipper locations from pixel coordinates to centimeters, allowing for standardized spatial analysis across different experimental setups.

Our analysis focuses on three main aspects of mouse behavior: 1). Distance calculation: We compute the Euclidean distance between the mouse’s nose and the sipper port for each frame. Utilized NumPy’s linalg.norm function. Any length greater than the width of the box was clipped to the width value. 2). Head direction determination: We compare the x value of the neck position against the x values of the body positions. The body positions were averaged then checked whether the x position of the neck was greater than the body average. If it was, we labeled the frame as facing right. 3). Turning identification: We identify turning behavior based on the sign changes in the difference in distances. We only identified significant turns where the change in direction was greater than 1.5 cm. We also only analyzed every second instead of the whole series to filter out noise.

To predict periods of engagement with the sipper port, we employ a multi-step filtering process: 1). Naive engagement identification: We initially classify frames as engaged if the nose is within 2 cm of the sipper port. 2). Anatomical feasibility filter: We remove frames where the nose is more than 3 cm away from the neck, as these represent anatomically infeasible postures. 3). Direction filter: We exclude frames where the mouse is not facing the sipper, based on our head direction calculation. 4). Stability filter: We remove frames where the neck is more than 4 cm from the sipper, indicating unstable nose position. 5). Temporal coherence filter: We filter out engagement periods shorter than 1 second, with a 0.5-second tolerance for brief disengagements, to focus on sustained engagement behaviors.

### Generalized linear model

#### Inputs

To analyze relationships between neural activity and behavior during cue periods, we developed a pipeline that processes and aligns multiple behavioral measurements. This pipeline requires four inputs: (1) SLEAP joint output files in H5 format, containing joint coordinates of shape (2, 30, n) where n is the number of frames; (2) engagement periods as timestamp pairs in Excel format; (3) lick events as timestamp pairs in Excel format; and (4) cue period timestamps in Excel format.

##### Data preprocessing

Our preprocessing involves three main stages. First, we convert all behavioral events into binary time series, where 1 indicates the presence of a behavior and 0 indicates its absence. Second, we extract additional behavioral states (walk, rear, groom, rest) using a pre-trained random forest classifier on the SLEAP joint data. Finally, we identify disengagement events, defined as the first frame following each engagement period.

##### Behavior alignment

We align all behavioral time series to cue onset, extracting data from 10 seconds pre-cue to 34 seconds post-cue. For each session, we aggregate behavior across trials by summing the binary arrays rather than averaging them. This summation approach creates a behavioral density measure that better corresponds to our calcium imaging data structure, which represents aggregate neural activity across the session rather than trial-level responses. Using density measures instead of trial-averaged behavior ensures that our predictor variables (behavioral measures) maintain the same temporal structure as our response variables (calcium signals). These density measures are then Z-scored to standardize their scale across sessions and behaviors.

##### Model fitting

We employ two levels of generalized linear model (GLM) analysis: 1). Cluster-level modeling: We fit GLMs using the normalized behavioral densities as predictors and mean cluster activity as the response variable. We use a Gaussian distribution with an identity link function. 2). Neuron-level modeling: Individual neuron activity serves as the response variable, with the same model structure as the cluster-level analysis. For both analyses, we extract and store model coefficients (β), their associated p-values, and model intercepts. Since we are using the identity link function, the β coefficients directly represent the strength and direction of the relationship between each behavior and neural activity.

##### Model validation

We used residual analysis to verify model assumptions.

### Quantification and Statistical Analysis

Unless otherwise noted, all data are expressed as mean ± SEM. Behavioral data were analyzed with GraphPad Prism 10.0 (GraphPad, La Jolla, CA). Two-tailed *student’s* t-test, one-way or two-way ANOVAs were used to analyze between-subjects designs. Holm-Sidak was used for *post-hoc* pairwise comparisons. The null hypothesis was rejected at the *p* < 0.05 level. Statistical significance was taken as \**p* < 0.05, \*\**p* < 0.01, \*\*\**p* < 0.001, \*\*\*\**p* < 0.0001 *n*.*s*. represents not significant.

### Materials Availability

This study did not generate new or unique reagents or other materials.

## Abbreviations

NAc: Nucleus Accumbens
PVT: Paraventricular Thalamus
eCB: endocannabinoid

## Data Availability

Source data are provided with this paper.

## Code Availability

Custom MATLAB analysis code was created to appropriately organize, process, and combine fiber photometry data with associated behavioral data. Analysis code for photometry from Figures 1, 2, 3, 4, and 5 will be made available online at https://www.github.com/BruchasLab.

## Acknowledgments

We thank the Molecular Genetics Resource Core for the Center in Neurobiology of Addiction, Pain, and Emotion and its director, Dr. Selena Schattauer, for generating the GRAB_eCB2.0_ and CRISPR viruses used in this manuscript. We thank Dr. Azra Suko for lab management and organization, as well Taylor Hobbs, Carina Pizzano, and Valerie Lau for colony management. Lastly, we thank the entire Bruchas lab as well as other members of the NAPE center at the University of Washington for resources and critical feedback.

## Author Contributions

D.J.M. performed conceptualization, methodology, investigation, formal analysis, fiber photometry analysis, 1-photon analysis, and writing of the original draft. A.E.E. designed and constructed the Linear Track, as well as wrote/adapted the code for subsequent deep learning classifications of behavior. G.S. wrote custom codes and analysis pipelines for Fourier/Rayleign analyses and GLM. E.F.S, R.O., S.H., and B.W. performed behavioral experiments, scored behavioral videos, and performed immunohistochemistry. S.C. wrote custom matlab codes for 1-photon analysis. A.Z. provided technical and experimental support. Y.L. designed and characterized GRAB_eCB2.0_. L.S.Z. designed the CRISPR viruses. N.S. provided funding for and aided with the conceptualization of the project. M.R.B. conceptualized, acquired funding for, and supervised the project, as well as assisted in writing the manuscript.

## Lead Contact

Further information and requests for resources and reagents should be directed to and will be fulfilled by the Lead Contact, Dr. Michael Bruchas.

## SUPPLEMENTAL FIGURES

**SUPPLEMENTAL FIGURE 1.**
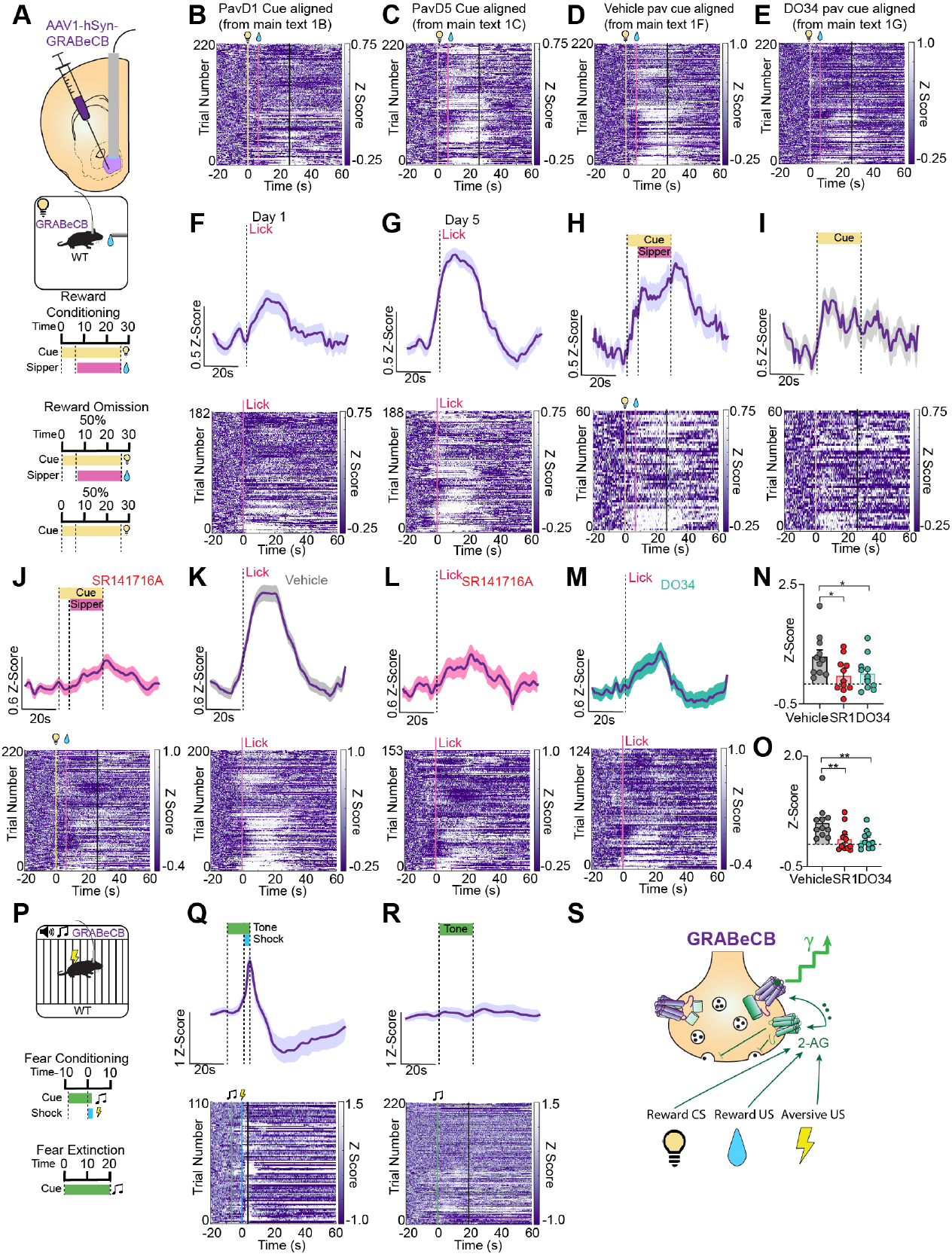
Endogenous cannabinoids are released in the NAc following rewarding and aversive stimuli. Schematic for fiber photometry recordings of GRAB_eCB2.0_ in the NAc during Pavlovian reward conditioning. (A). Heatmaps of GRAB_eCB2.0_ in the NAc corresponding to the traces in main text figure 1B, 1C, 1F, and 1G, respectively (B-E). Photometry trace and heatmap of lick-aligned GRAB_eCB2.0_ signal on day 1 (F) and day 5 (G) of reward conditioning. Photometry trace and heatmap of lick-aligned GRAB_eCB2.0_ signal during rewarded (H) and non-rewarded (I) trials in the reward omission assay. Photometry trace and heatmap of cue-aligned GRAB_eCB2.0_ signal in SR141716A treated animals (J). Photometry trace and heatmap of lick-aligned GRAB_eCB2.0_ signal in vehicle (K), SR141716A (L), and DO34 (M) treated animals. Cue-aligned Z-scores showing attenuation of lick-aligned GRAB_eCB2.0_ signal in SR141716A (p=0.0397) and DO34 (p=0.0397) treated animals compared to vehicle (n=11) (N). Comparison of cue-aligned GRAB_eCB2.0_ signal in SR141716A (p=0.0083) and DO34 (p=0.0083) treated animals (N=11) (O). Schematic for fiber photometry recordings of GRAB_eCB2.0_ in the NAc during Pavlovian reward conditioning (P). Photometry trace and heatmap of shock-aligned GRAB_eCB2.0_ signal during fear conditioning (Q). Photometry trace and heatmap of cue-aligned GRAB_eCB2.0_ signal during fear extinction (R). Schematic depicting stimulus-elicited eCB release in the NAc (S). All error bars represent ± SEM; N represents number of mice. p values reported from one-way ANOVA \**p* < 0.05, \*\**p* < 0.01 (N,O).

**SUPPLEMENTAL FIGURE 2.**
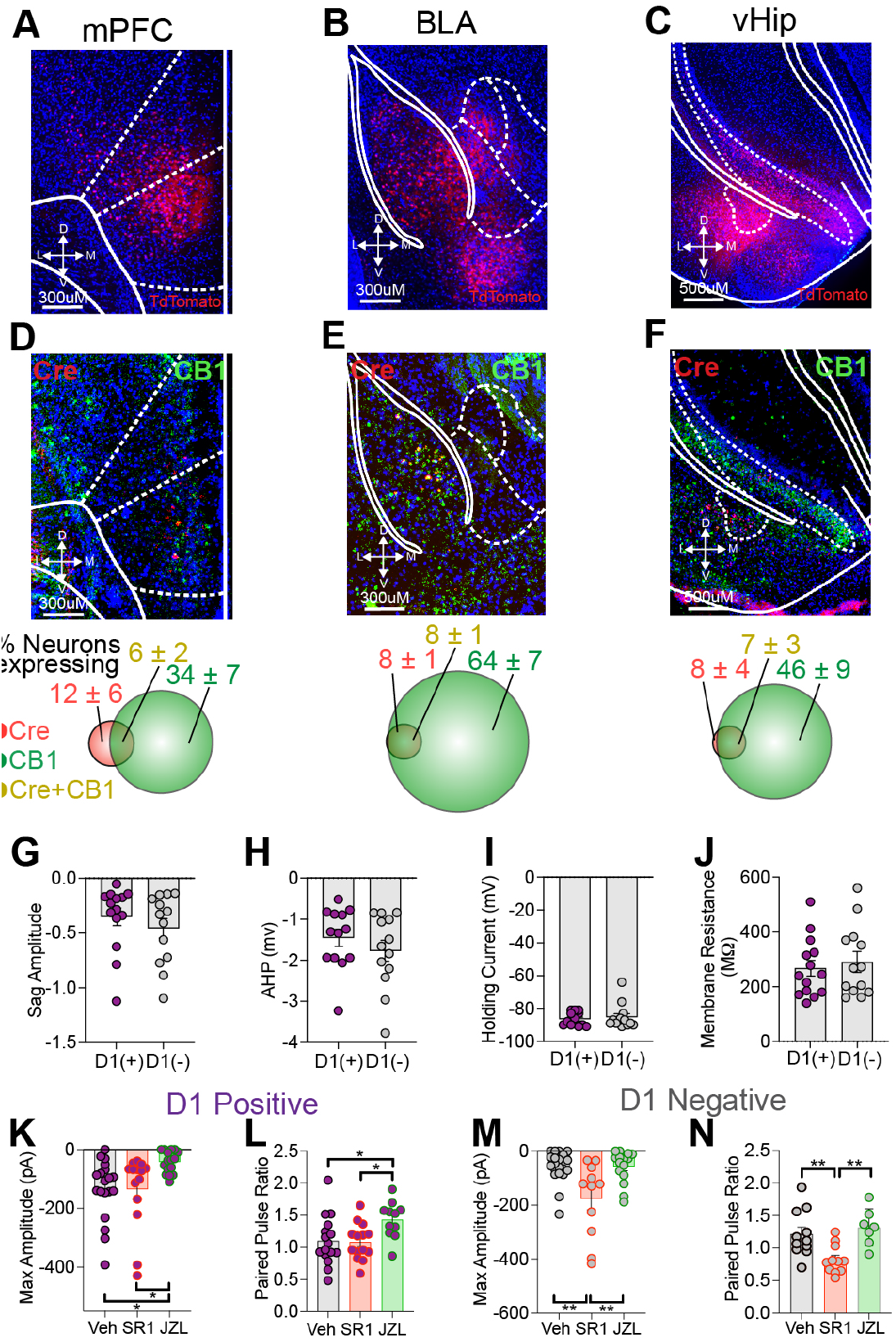
Histological analysis of CB_1_R expressing inputs to the NAc and electrophysiological characterization of excitatory aPVT^NTS^-NAc projections. TdTomato positive NAc projecting neurons in the medial prefrontal cortex (mPFC) (A) basolateral amygdala (BLA) (B) and ventral hippocampus (vHip) (C). RNAscope image and quantification of Cre and GRAB_eCB2.0_ transcript levels in the mPFC (D), BLA (E), and vHip (F). Electrophysiological comparison of sag current amplitude (p=0.3871) (G) afterhyperpolzarization voltage (p=0.3404) (H), holding current (p=0.6077) (I), and membrane resistance (p=0.6166) (J) between D_1_R(+) (n=14) and D_1_R(-) (n=13) neurons. 1JZL184 treatment significantly reduces the maximum oEPSC amplitude (p=0.0193) (K) and increases the Paired Pulse Ratio (p=0.0355) (L) in D_1_R(+) neurons compared to vehicle treated slices (Veh: n=19, SR141716A: n=14, JZL184: n=16). 10SR141716A significantly increases the maximum oEPSC amplitude (p=0.0016) (M) and reduces the Paired Pulse Ratio (p=0.0092) (N) in D_1_R(-) neurons (Veh: n=20, SR141716A: n=11, JZL184: n=15). All error bars represent ± SEM; n represent number of neurons. p values reported from two-tailed unpaired t-test (G,H,I,J) and one-way ANOVA \**p* < 0.05, \*\**p* < 0.01 (K,L,M,N).

**SUPPLEMENTAL FIGURE 3.**
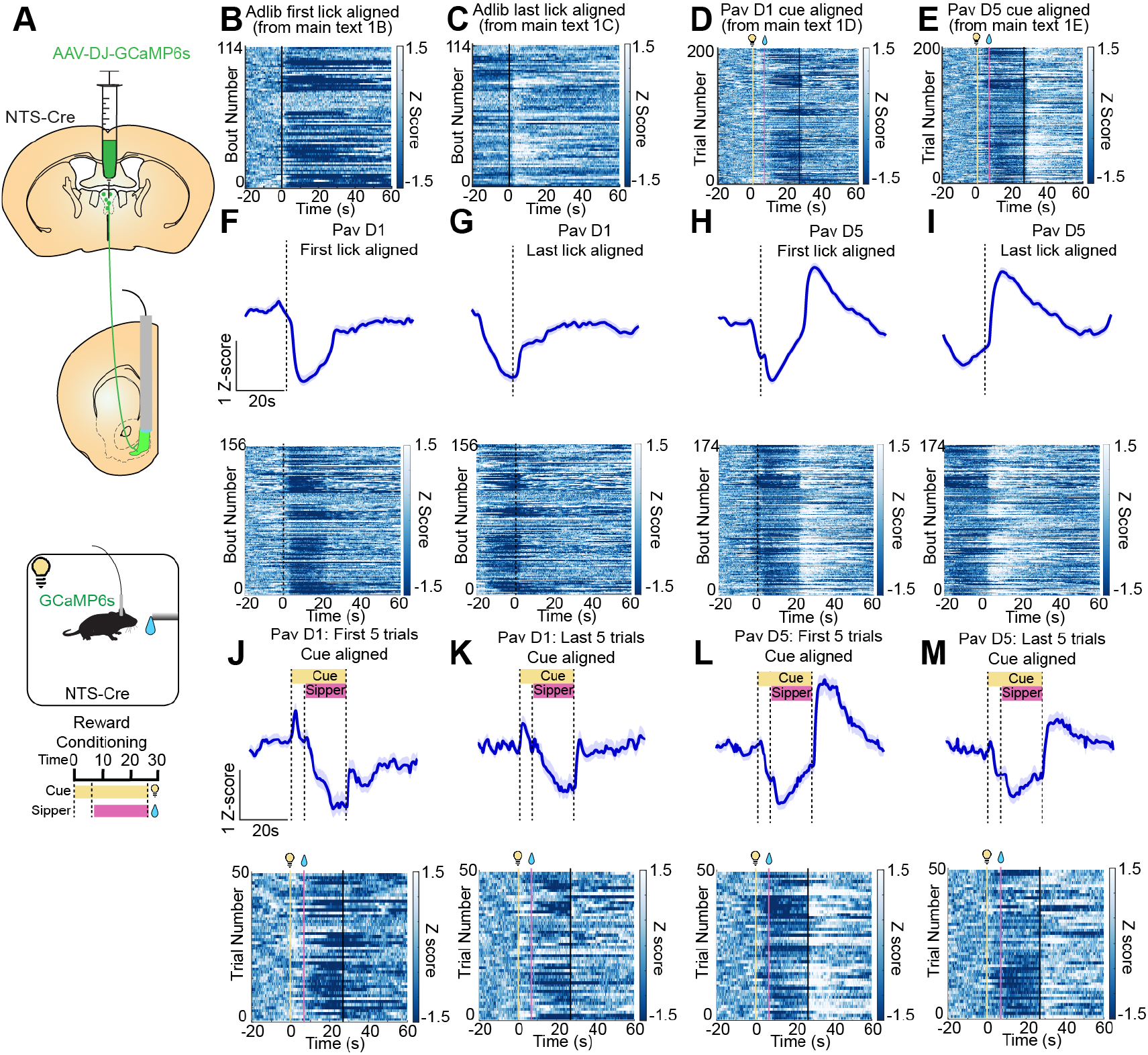
Figure S3: Consumption onset and offset-aligned aPVT^NTS^-NAc GCaMP6s dynamics. Schematic for fiber photometry recordings from GCaMP6s expressing aPVT^NTS^-NAc terminals during Pavlovian reward conditioning (A). Heatmaps of GCaMP6s signal from aPVT^NTS^-NAc terminals corresponding to the traces in main text figure 1B, 1C, 1D, and 1E, respectively (B-E). Photometry trace and heatmap of GCaMP6s signal aligned to first lick (F) and last lick (G) on day 1 of reward conditioning. Photometry trace and heatmap of GCaMP6s signal aligned to first lick (H) and last lick (I) on day 5 of reward conditioning. Photometry trace and heatmap of GCaMP6s signal aligned to cue during the first 5 (J) and last 5 (K) trials on day 1 of reward conditioning. Photometry trace and heatmap of GCaMP6s signal aligned to cue during the first 5 (L) and last 5 (M) trials on day 5 of reward conditioning.

**SUPPLEMENTAL FIGURE 4.**
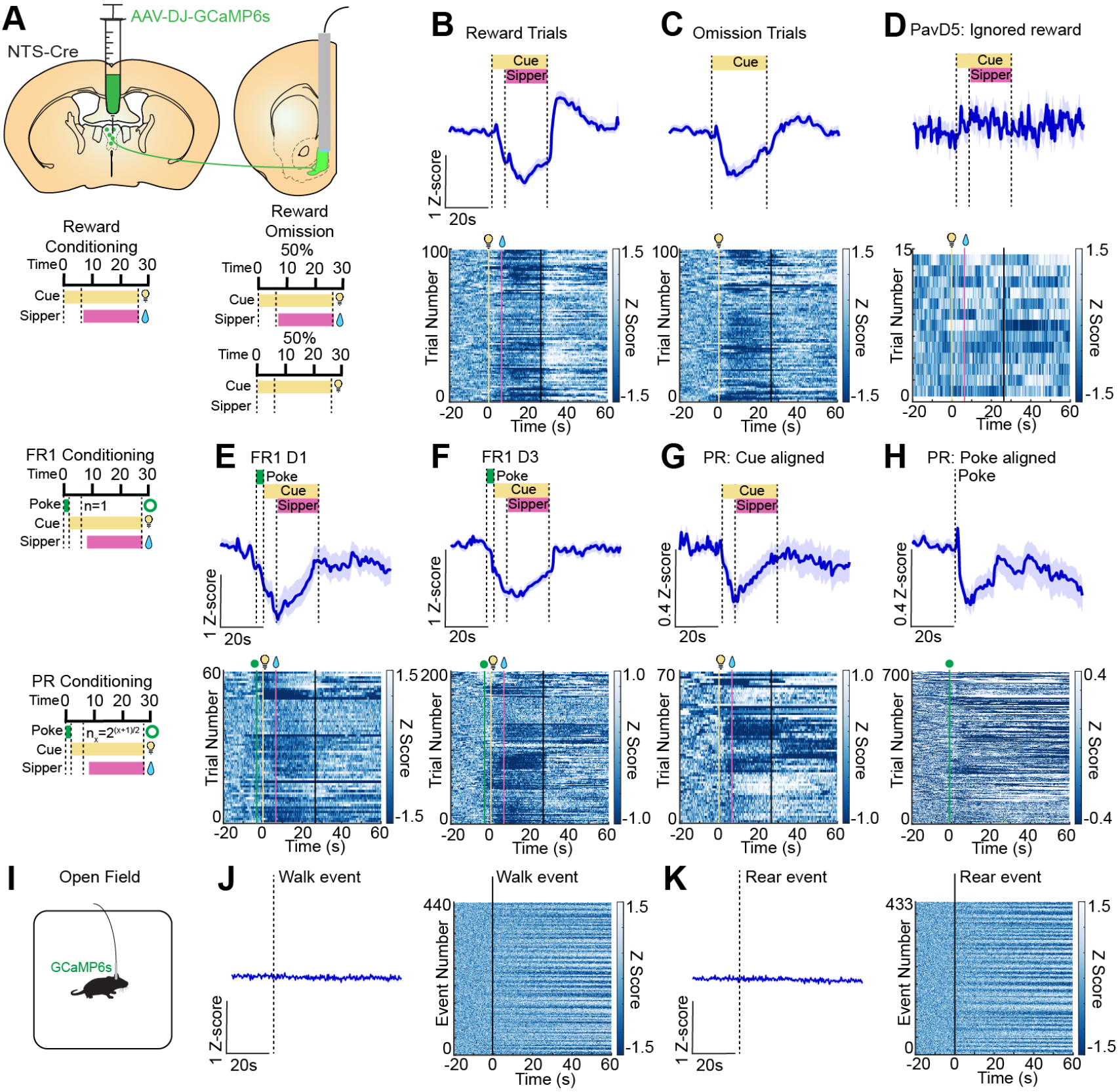
Inhibition of aPVT^NTS^-NAc terminals is not dependent on reward receipt and is not associated with locomotor activity. Schematic for fiber photometry recordings from GCaMP6s expressing aPVT^NTS^-NAc terminals during Pavlovian reward conditioning, reward omission, Fixed Ratio 1 (FR1) training, and Progressive Ratio (PR) training (A). Photometry trace and heatmap of GCaMP6s signal during rewarded (B) and non-rewarded (C) trials during the reward omission test, demonstrating aPVT^NTS^-NAc terminal inhibition in the absence of reward receipt. Photometry trace and heatmap of GCaMP6s signal on day 5 of reward conditioning during ignored cue/reward trials (no reward engagement), demonstrating a lack of inhibition when the cue/reward is ignored (D). Photometry trace and heatmap of GCaMP6s signal aligned to cue on day 1 (E) and day 3 (F) of FR1 conditioning, demonstrating inhibition during active operant responding for reward. Photometry trace and heatmap of GCaMP6s signal aligned to cue (G) and nose poke (H) during PR conditioning. Schematic for fiber photometry recordings of GCaMP6s from aPVT^NTS^-NAc terminals during the open field assay (I). Photometry trace and heatmap of GCaMP6s signal aligned to walk (J) and rear (K) events during the open field assay.

**SUPPLEMENTAL FIGURE 5.**
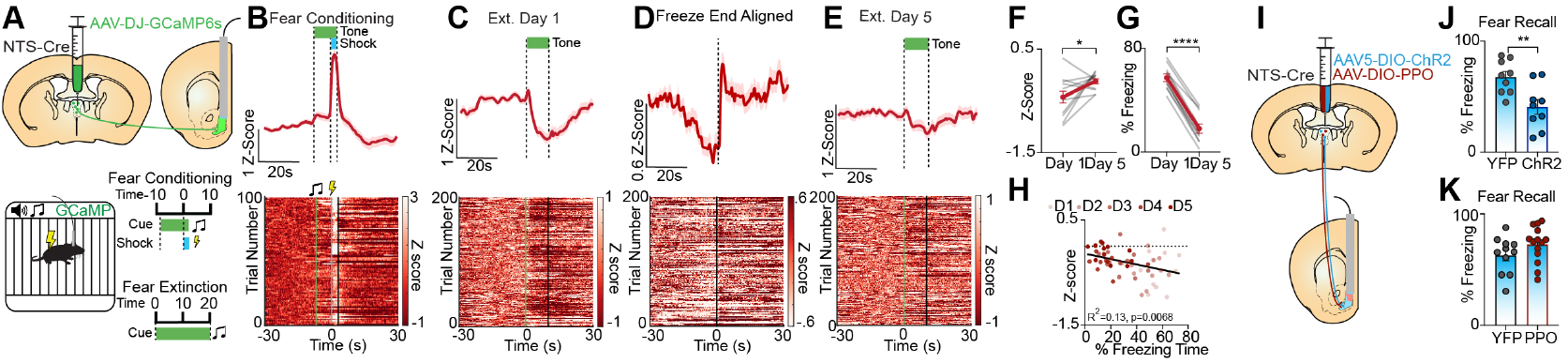
aPVT^NTS^-NAc terminal activity is negatively correlated with engagement in defensive freezing behaviors. Schematic for fiber photometry recordings from GCaMP6s expressing aPVT^NTS^-NAc terminals during Pavlovian fear conditioning and extinction (A). Photometry trace and heatmap of GCaMP6s signal aligned foot shock during fear conditioning (B). Photometry trace and heatmap of GCaMP6s signal aligned to cue presentation (C) and freezing cessation (D) on day 1 of fear extinction. Photometry trace and heatmap of GCaMP6s signal aligned to cue presentation on day 5 of fear extinction (E). Comparison between extinction day 1 and day 5 of average Z-score (N=11, p=0.0318) (F) and freezing time (N=11, p=<0.0001) (G) during the 20 second cue presentation. Correlation between freezing time and Z-score across 5 days of extinction (N=11, R2=0.13, p=0.0068) (H). Schematic for optogenetic manipulation of aPVT^NTS^-NAc terminals using DIO-ChR2 and DIO-PPO (I). 20hz optical stimulation ChR2 (N=9) expressing aPVT^NTS^-NAc terminals during the 20 second cue presentation decreases freezing time compared to eYFP (N=9) controls (p=0.0053) (J). 10hz optical stimulation of PPO (N=11) expression aPVT^NTS^-NAc terminals does not alter freezing behavior compared to eYFP (N=14) controls (p=0.1470) (K). All error bars represent ± SEM; N represent number of mice. p values reported from paired two-tailed t-test (F,G), linear regression (H), and unpaired two-tailed t-test (J,K) \**p* < 0.05, \*\**p* < 0.01, \*\*\**p* < 0.001, \*\*\*\**p* < 0.0001.

**SUPPLEMENTAL FIGURE 6.**
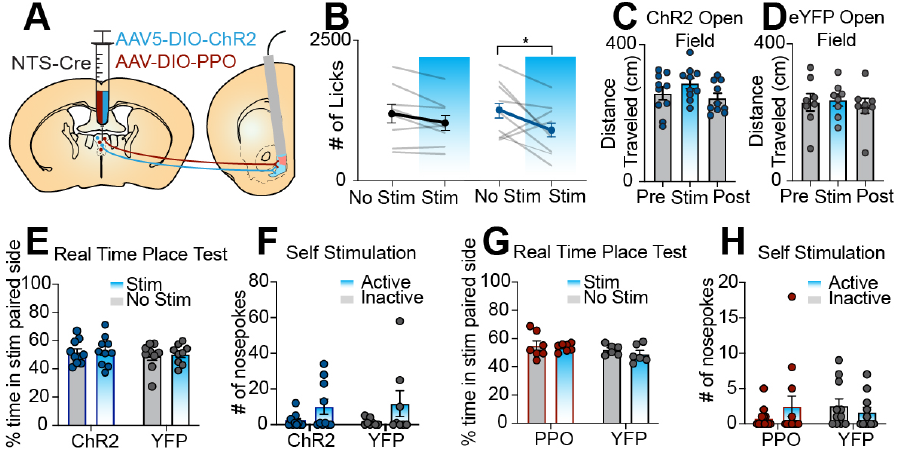
Neither activation nor inhibition of aPVT^NTS^-NAc terminals alters locomotion, drives real-time place preference (RTPP), or drive intracranial self-stimulation (ICSS) Schematic for optogenetic manipulation of aPVT-NAc terminals using DIO-ChR2 and DIO-PPO (A). 20hz photo-stimulation of ChR2 expressing aPVT^NTS^-NAc terminals during cue presentation reduces engagement in consummatory behaviors (ChR2: N=9, eYFP: N=8), p=0.0427) (B). Neither 20hz photo-stimulation (2 mins off, 2 mins on, 2 mins off) of ChR2 expressing aPVT^NTS^-NAc terminals (N=10, p=0.3655) (C) nor eYFP expressing aPVT^NTS^-NAc terminals (N=8, p=0.8947) (D) alters locomotor behavior. 20hz photo-activation of aPVT^NTS^-NAc terminals does not support RTPP (ChR2: N=10 p=0.8141, eYFP: N=9 p=0.7191) (E). 20hz photo-activation (2 second burst per nose poke) of aPVT^NTS^-NAc terminals does not support ICSS (ChR2: N=10 p=0.0968, eYFP: N=8, p=0.1670) (F). 10hz photo-inhibition of aPVT^NTS^-NAc terminals does not support RTPP (PPO: N=7 p=0.7492, eYFP: N=6 p=0.3216) (G). 10hz photo-inhibition of aPVT^NTS^-NAc terminals does not support ICSS (PPO: N=13 p=0.2565, eYFP: N-=11 p=0.4527) (H). All error bars represent ± SEM; N represent number of mice. p values reported from paired two-tailed t-test (B,E,F,G,H), and one-way ANOVA (C,D). \**p* < 0.05.

**SUPPLEMENTAL FIGURE 7.**
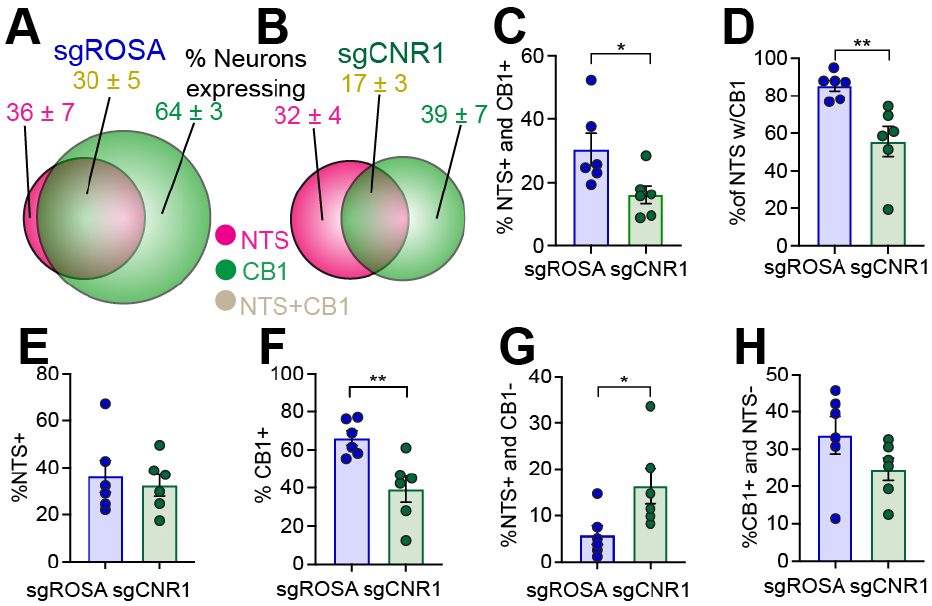
Characterization of AAV1-FLEX-sgCNR1 deletion of CB_1_R from aPVT NTS neurons. Diagram showing % of neurons expressing NTS, CB_1_R, or NTS and CB_1_R in the aPVT from NTS-Cre mice injected with sgROSA (N=6) (A) or sgCNR1 (N=6) (B). % of total neurons expressing NTS and CB_1_R (p=0.0333) (C). % of NTS neurons expressing CB_1_R (p=0.0055) (D). % of total neurons expressing NTS (p=0.6507) (E). % of total neurons expressing CB_1_R (p=0.0064) (F). % of total neurons that are NTS positive and CB_1_R negative (p=0.0354) (G). % of total neurons that are CB_1_R positive and NTS negative (p=0.1512) (H). All error bars represent ± SEM; N represent number of mice. p values reported from unpaired two-tailed t-test (C,D,E,F,G,H) \**p* < 0.05, \*\**p* < 0.01.

**SUPPLEMENTAL FIGURE 8.**
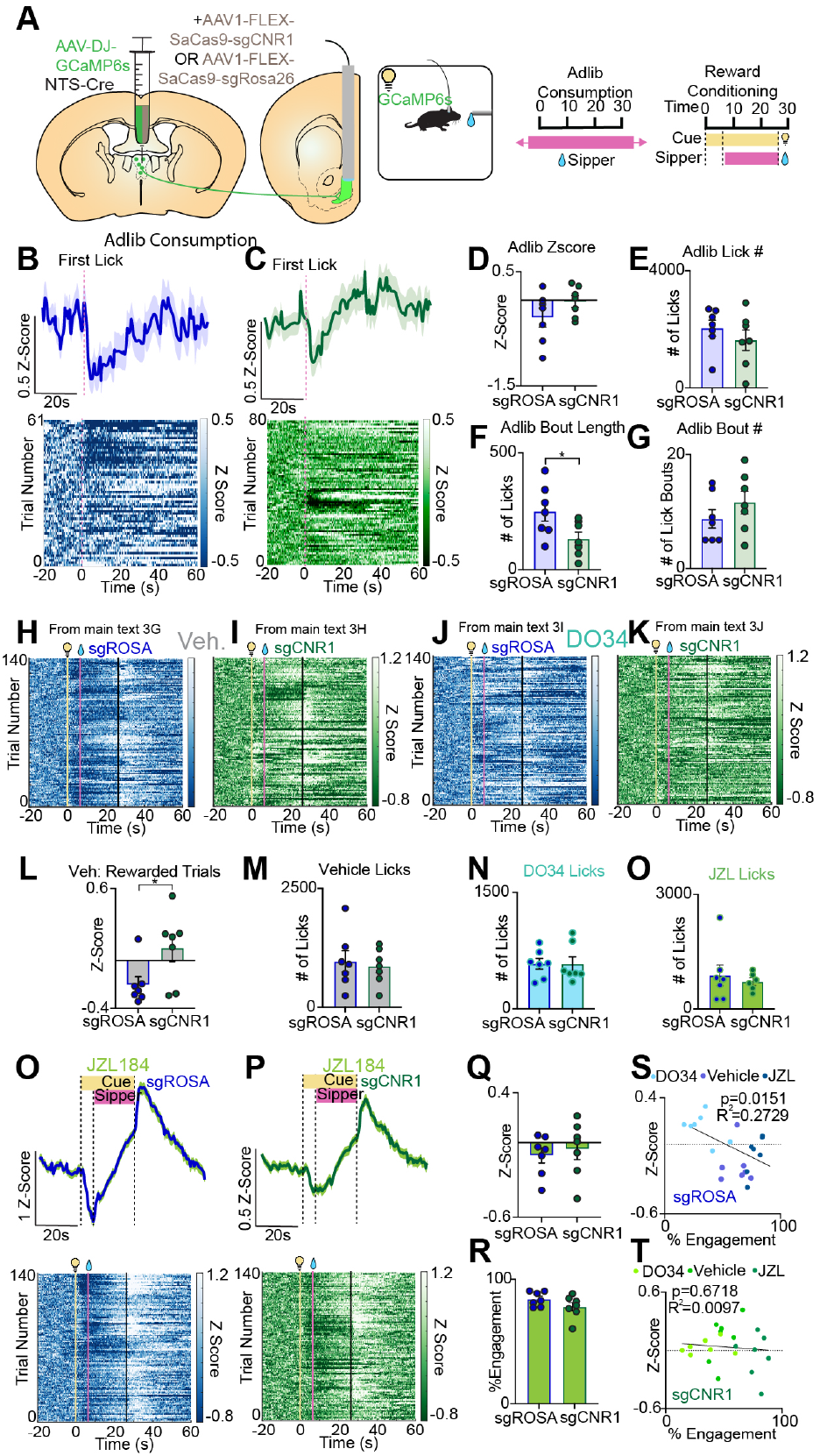
Effect of AAV1-FLEX-sgCNR1 deletion of CB_1_R from aPVT NTS neurons during reward consumption and Pavlovian reward conditioning. Schematic for CB_1_R deletion from the aPVT and photometry recordings of GCaMP6s aPVT^NTS^-NAc terminals during adlib sucrose consumption and Pavlovian reward conditioning (A). Heatmaps of GCaMP6s signal from aPVT^NTS^-NAc terminals corresponding to the traces in main text figure 3G, 3H, 3I, and 3J, respectively (B-E). Photometry trace and heatmap of GCaMP6s signal aligned first lick during adlib sucrose consumption from sgROSA (N=7) (F) and sgCNR1 (N=7)(G) expressing mice. Comparison of first lick-aligned photometry Z-score between sgROSA and sgCNR1 expressing mice (p=0.1741) (H) during adlib sucrose consumption. Comparison of total lick number (p=0.3705) (I), bout length (p=0.0393) (J), and bout number (p=0.2828) (K) between sgROSA and sgCNR1 expressing mice during adlib sucrose consumption. Comparison of photometry Z-score during rewarded trials in vehicle treated sgROSA and sgCNR1 expressingmice during Pavlovian reward conditioning (p=0.0373) (L). Comparison of number of licks in vehicle (p=0.7029) (I), DO34 (p=0.9883) (M), and JZL184 (p=0.6107) (N) treated sgROSA and sgCNR1 expressing mice during Pavlovian reward conditioning. Photometry trace and heatmap of GCaMP6s signal aligned first lick during Pavlovian reward conditioning from JZL184 treated sgROSA (O) and sgCNR1 (P) expressing mice. Comparison of photometry Z-score (0-26 seconds, p=0.6359) (Q) and % sipper port engagement time (0-26 seconds, p=0.1569) (R) during Pavlovian reward condition from JZL184 treated sgROSA and sgCNR1 expressing animals. Linear regression of % sipper port engagement time and photometry Z-score in sgROSA expressing animals (R2=0.2729, p=0.0151) (S). Linear regression of % sipper port engagement time and photometry Z-score in sgCNR1 expressing animals (R2=0.0097, p=0.6718) (T). All error bars represent ± SEM; N represent number of mice. p values reported from unpaired two-tailed t-test (H, I, J, K, L, M, N, O, Q, R) or linear regression (S,T) \**p* < 0.05.

**SUPPLEMENTAL FIGURE 9.**
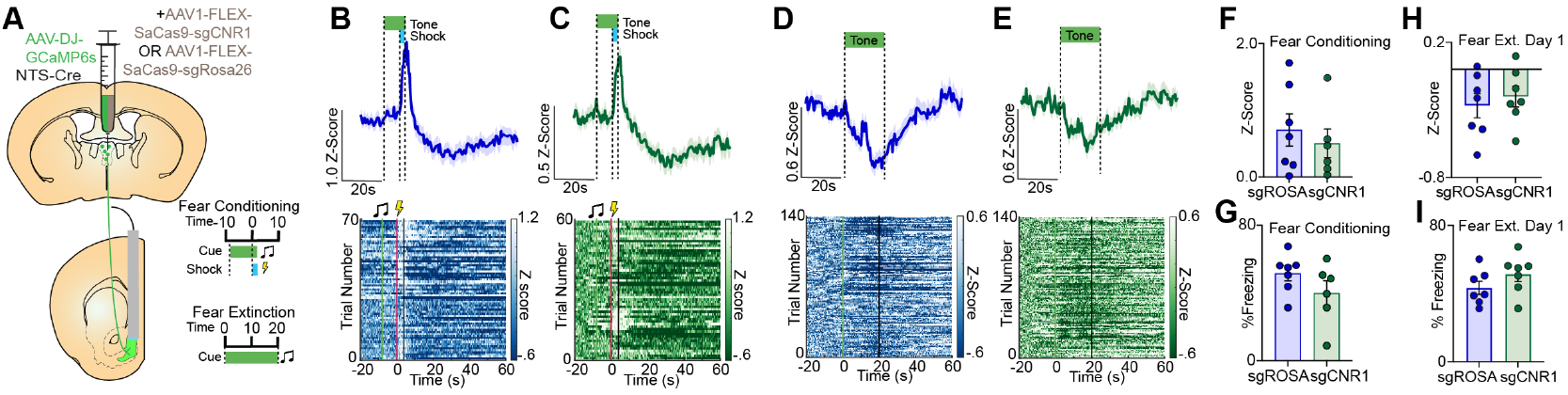
AAV1-FLEX-sgCNR1 deletion of CB_1_R from aPVT NTS neurons does not affect fear conditioning or extinction. Schematic for CB_1_R deletion from the aPVT and photometry recordings of GCaMP6s aPVT^NTS^-NAc terminals during Pavlovian fear conditioning and extinction (A). Photometry trace and heatmap of GCaMP6s signal aligned to foot shock from sgROSA (N=7) (B) and sgCNR1 (N=7)(C) expressing mice during Pavlovian fear conditioning. Photometry trace and heatmap of GCaMP6s signal aligned to cue presentation from sgROSA (N=7) (D) and sgCNR1 (N=7)(E) injected mice during fear extinction. Comparison of freezing behavior between sgROSA and sgCNR1 injected animals during fear conditioning (p=0.1851) (F) and day 1 of fear extinction (p=0.1873) (G). Comparison of photometry Z-score between sgROSA and sgCNR1 injected animals during fear conditioning (p=0.7027) (H) and day 1 of fear extinction (p=0.5777) (I). Schematic for photometry recordings of GRABeCB2.0 or GRAB_eCB2.0MUT_ during fear conditioning and fear extinction (J). Photometry trace and heatmap of GRABeCB2.0 (n=8) or GRAB_eCB2.0MUT_ (n=4) signal during fear conditioning (K) or fear extinction (L). All error bars represent ± SEM; N represent number of mice. p values reported from unpaired two-tailed t-test (F,G,H, I) \**p* < 0.05, \*\**p* < 0.01.

**SUPPLEMENTAL FIGURE 10.**
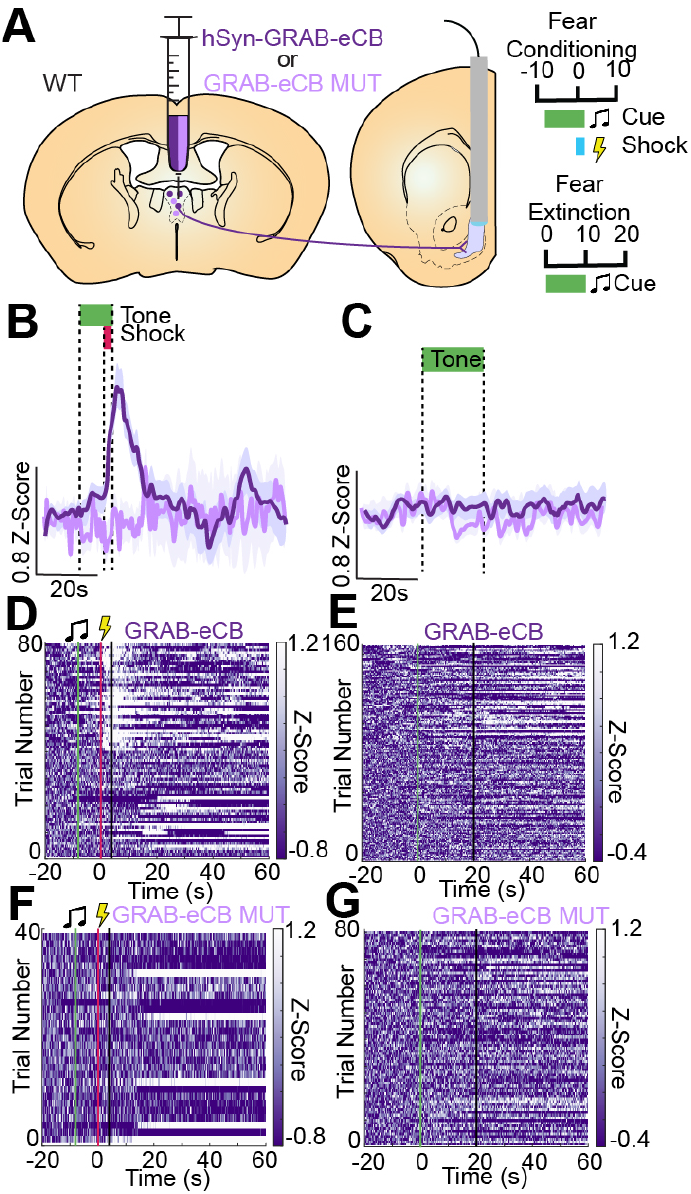
Foot shock induces eCB release and binding to aPVT terminals in the NAc. Schematic for photometry recordings of GRAB_eCB2.0_ or GRAB_eCB2.0MUT_ during fear conditioning and fear extinction (A). Photometry trace and heatmap of GRAB_eCB2.0_ (n=8) or GRAB_eCB2.0MUT_ (n=4) signal during fear conditioning (B,D,F) or fear extinction (C,E,G)

**SUPPLEMENTAL FIGURE 11.**
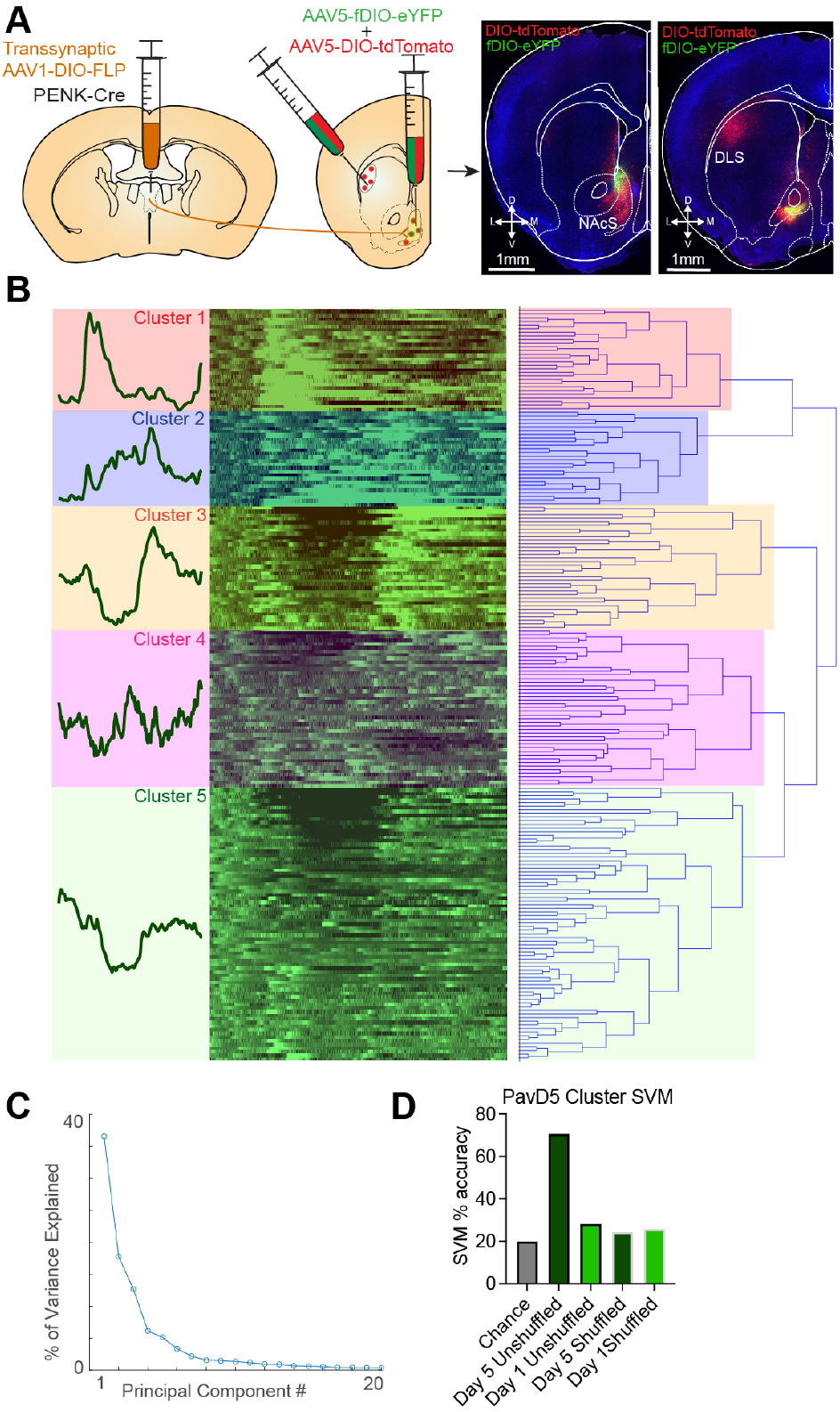
Characterization of transsynaptic labeling and cluster classification. Schematic for validation of transsynaptic labeling approach. Transsynaptic AAV1-DIO-FLP was injected into the aPVT, AAV5-DIO-tdTomato was injected into the dorsolateral striatum (DLS), and a cocktail of AAV5-DIO-tdTomato and AAV5-fDIO-eYFP was injected in the NAc of PENK-Cre mice. The DLS, which expresses PENK but does not receive aPVT input, showed expression of AAV5-DIO-tdTomato but not AAV5-DIO-eYFP. The NAc, which expresses PENK and does receives aPVT input, showed expression of both AAV5-DIO-tdTomato and AAV5-DIO-eYFP (A). Hierarchical clustering of 206 tracked aPVT-NAc^Penk^ neurons based on Principal Component Analysis (PCA) of the activity of each neuron in the 60 second window following cue-onset. In order to be considered a cluster, the number of neurons in that cluster had to represent at least 10% of the total population of neurons (>21 neurons) based on our a priori criteria (B). Percent of variance explained by the top 20 PCs (C). Support Vector Machine decoding of cluster identity, trained on PavD5 cluster activity (D)

**SUPPLEMENTAL FIGURE 12.**
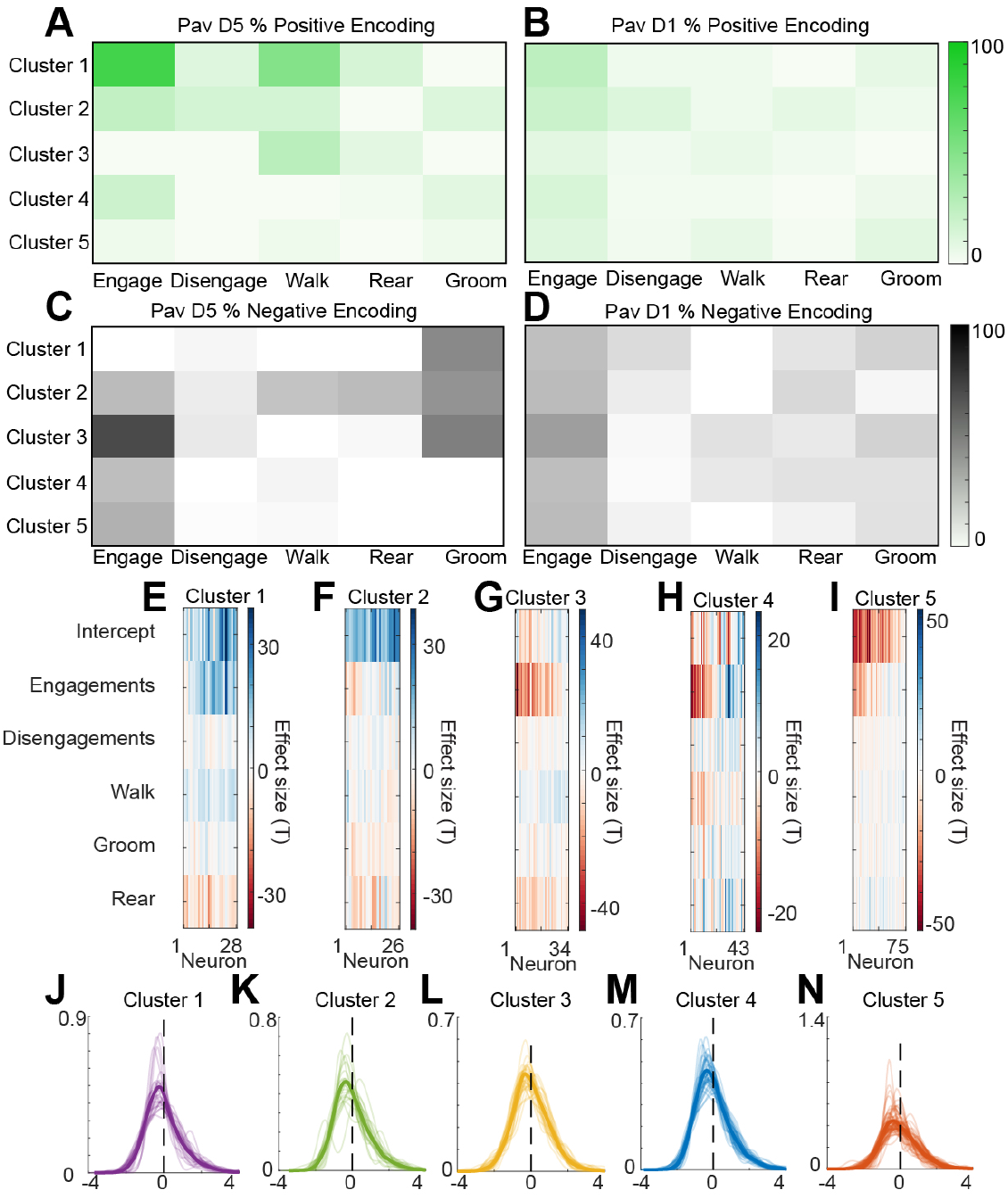
Validation of generalized linear model. Percent of neurons within each cluster encoding each behavioral state, with a beta coefficient threshold of +/-0.5 (A-D). Contribution of effect sizes for each neuron in each cluster to the generalized linear model (T values) (E-I). Raw residual distributions from the generalized linear model for each neuron in each cluster (J-N).

**SUPPLEMENTAL FIGURE 13.**
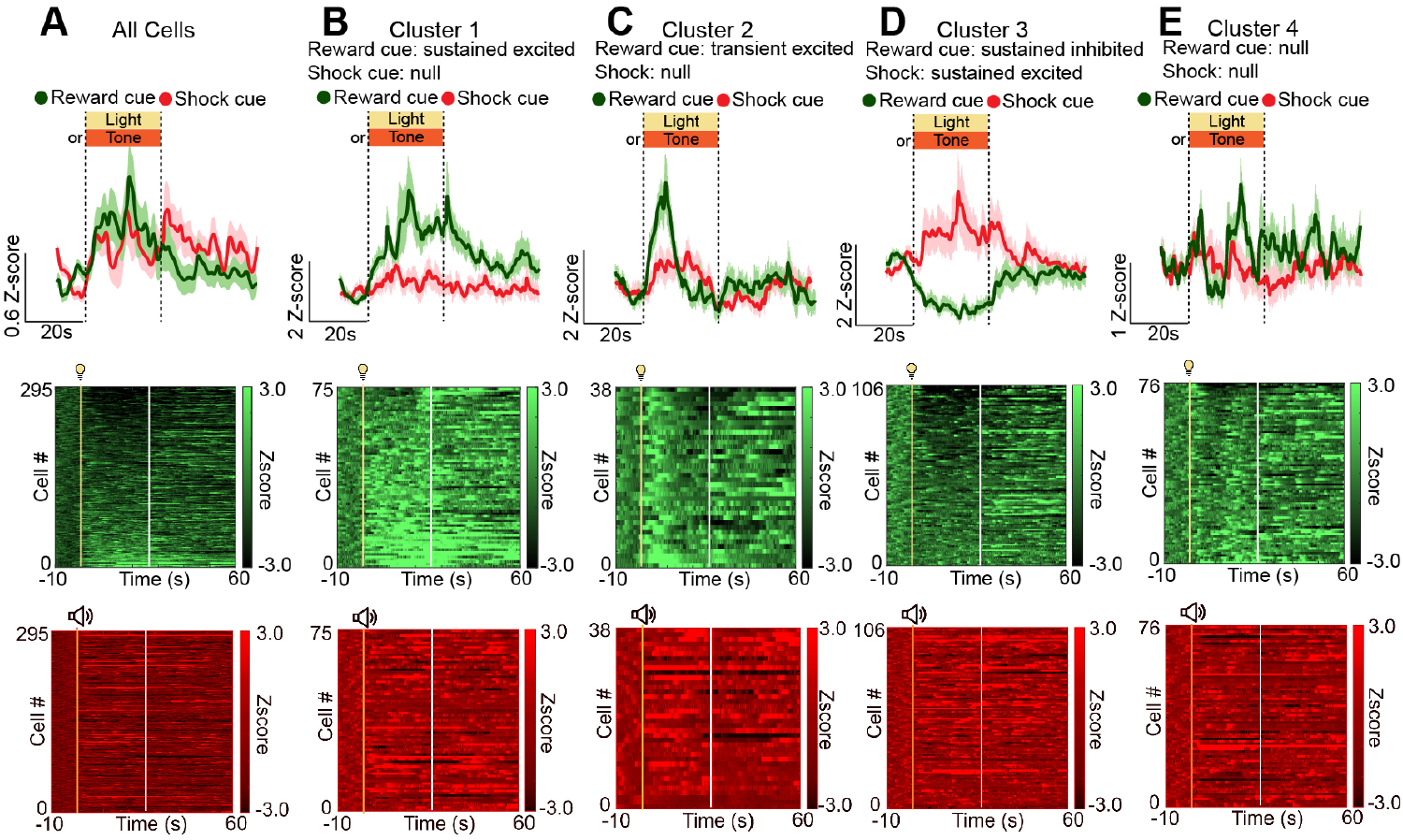
Unique encoding of reward and aversion predictive cues during a cue-discrimination task. Average reward predictive cue-aligned trace and heatmaps for 206 neurons as well as the neuron-matched trace and heatmap for the shock-predictive cue (N=10) (A). Traces and heatmaps from 4 unique reward predictive cue-aligned clusters of aPVT-NAc^Penk^ neurons, as well as their neuron-matched traces and heatmaps for the shock-predictive cue (B-E).

**SUPPLEMENTAL FIGURE 14.**
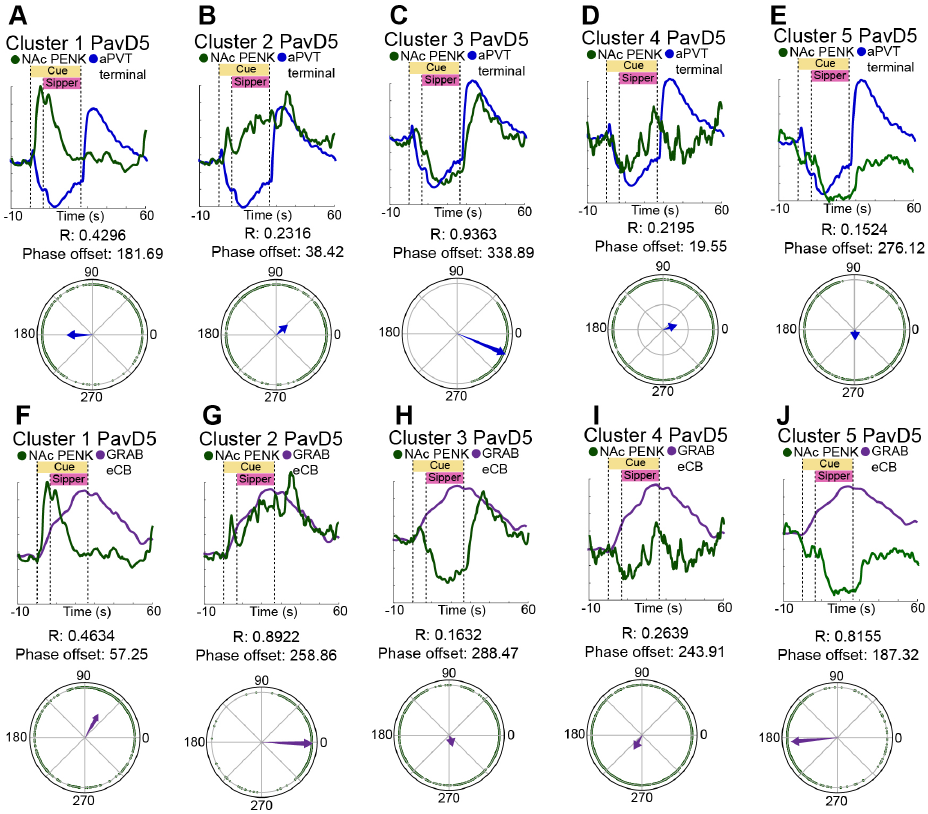
Hilbert and Rayleigh analysis of cluster entrainment to aPVT^NTS^-NAc GCaMP6s and aPVT-NAc GRAB_eCB2.0_ signal. Overlay of Pavlovian reward conditioning day 5 aPVT-NAc^Penk^ cluster GCaMP6s traces with aPVT^NTS^-NAc terminal GCaMP6s traces and Rayleigh plots from cluster 1 (A), 2 (B), 3 (C), 4 (D), and 5 (E). Overlay of Pavlovian reward conditioning day 5 aPVT-NAc^Penk^ cluster GCaMP6s traces with aPVT-NAc terminal GRAB_eCB2.0_ traces and Rayleigh plots from cluster 1 (F), 2 (G), 3 (H), 4 (I), and 5 (J).

